# Mitochondria-Lysosome Crosstalk Shapes Metabolic Transition in Neonatal Enterocytes

**DOI:** 10.1101/2025.09.09.674148

**Authors:** Gonzalo Herranz, Diego Alonso-Larre, Tamara González, Laura Akintche, Alejandra Ramos-Manzano, Marta Iborra-Pernichi, María Velasco de la Esperanza, Covadonga Díaz-Díaz, Ian G Ganley, Patricia Boya, Sara Cogliati, Nuria Martínez-Martín, Fernando Martín-Belmonte

## Abstract

The neonatal gastrointestinal tract mediates nutrient absorption and the establishment of immune tolerance to commensal microbiota. In early life, lysosome-rich enterocytes (LREs) in the ileum are necessary for the intracellular digestion of maternal milk proteins. However, the molecular mechanisms sustaining their function remain incompletely characterized. Here, we demonstrate that LRE mitochondrial homeostasis and autophagic capacity are critical for efficient nutrient uptake and maintenance of their specialized identity, as disruption of either process leads to premature differentiation into post-weaning enterocytes (PECs) with diminished endolysosomal and metabolic activity. Transcriptomic profiling further revealed that neonatal LREs exhibit a distinctive antioxidant signature, which preserves redox balance and safeguards the expression of the transcriptional regulators MAFB and BLIMP1, both central repressors of the neonatal-to-adult metabolic transition. These findings establish the mitochondria–lysosome axis as a key determinant of LRE function and neonatal metabolic programming. They also provide a mechanistic framework for understanding how organelle dysfunction and redox imbalance may contribute to early-life malnutrition syndromes, such as Kwashiorkor, and suggest therapeutic strategies aimed at preserving mitochondrial and lysosomal integrity.

## INTRODUCTION

The small intestine is a specialized section of the gastrointestinal tract (GIT) engaged in nutrient digestion and absorption, while simultaneously acting as a barrier against harmful substances. It consists of a variety of cell types, with epithelial and immune cells playing key roles in maintaining intestinal function. Through their interactions with one another and the gut microbiota, these cells help create a finely balanced microenvironment that supports both nutritional and immunological functions^1,2^. The main absorptive epithelial cells in the intestine are called enterocytes. They are the most numerous cells in the intestinal epithelium and, along with secretory epithelial cells (including Paneth, goblet, and enteroendocrine cells), originate from stem cells of the intestinal crypts^1^.

Dietary protein is a macronutrient essential for growth and overall organismal homeostasis. In weaned mammals, protein digestion takes place in the lumen of GIT, where gastric pepsin and intestinal proteases sequentially break down proteins into oligopeptides and free amino acids, which are then absorbed by enterocytes via specific transporters^3–5^.

However, in the intestinal lumen of suckling mammals and stomachless fishes, protease activity remains low and protein digestion is incomplete^3–5^. For this reason, their posterior-mid small intestine contains a population of lysosome-rich enterocytes (LREs), formerly referred to as vacuolated enterocytes^6–8^. These cells possess a highly developed endolysosomal system, including a prominent lysosomal vacuole (LV), which is capable of internalising and metabolising intact proteins via non-specific fluid-phase processes (pinocytosis) and receptor-mediated endocytosis and membrane adsorption^5,7,9,10^. Moreover, LREs facilitate the transcytosis of antigens, maternal immunoglobulins, and antibodies across the epithelial barrier while preserving their biological activity, contributing to the immune system development^5,11^. Recent studies have highlighted the nutritional importance of LREs during the neonatal period. Key endolysosomal proteins, whose absence leads to developmental defects, have been identified, including Mucolipins 1 and 3^12^, Pllp^8^, Mamdc4 (also known as endotubin)^13^ and Dab2^7^. Despite these findings, the molecular mechanisms that regulate endocytic and lysosomal pathways in LREs are still not fully understood.

Autophagy is a conserved lysosome-dependent catabolic pathway that recycles intracellular components and nutrients. This process involves the sequestration of cytoplasmic material into double-membraned autophagosomes, which subsequently fuse with lysosomes for degradation and breakdown. The lipidation of LC3, a key step in autophagosome formation, is mediated by the ATG12–ATG5–ATG16L1 complex^14,15^. Mutations in components of this complex have been linked to intestinal pathologies^16,17^, and *Atg5*-deficient neonates exhibit increased neonatal mortality under fasting conditions, underscoring the essential role of autophagy during early postnatal life^18^. Despite this, the homeostatic role of autophagy throughout the entire neonatal period, particularly in LRE biology, remains unclear.

Autophagy plays a crucial role in cellular metabolism and differentiation, primarily through the selective elimination of mitochondria (mitophagy), thereby contributing to the control of reactive oxygen species (ROS) levels^19,20^. Beyond their role in energy production, mitochondria and ROS also function as key regulators of cell signalling and differentiation^21–23^. In the intestinal epithelium, a gradient of increasing mitochondrial mass, activity, and ROS levels has been described along the crypt–villus axis, correlating with higher degrees of cellular differentiation^24^. However, how mitochondrial homeostasis and ROS dynamics influence the developmental remodelling of enterocytes remains poorly understood. In particular, their role in regulating the metabolic and functional transition of neonatal LREs into post-weaning enterocytes (PECs) has not yet been explored.

This study reveals that the mitochondria–lysosome axis is essential for the function and maturation of LREs. We show that LREs rely on high autophagic and mitophagic activity to sustain nutrient absorption and lysosomal integrity, and that disruption of either autophagy (*Atg16*cKO) or mitochondrial homeostasis (*Tfam*cKO) leads to impaired endolysosomal function, oxidative stress, and premature weaning transition through the repression of the transcription factors BLIMP1 and MAFB. These findings establish organelle homeostasis as a critical determinant of neonatal enterocyte identity and provide mechanistic insights into how its disruption contributes to childhood malnutrition.

## RESULTS

### Lysosome-rich enterocytes (LREs) display heightened basal autophagy levels essential for absorptive function

Unlike adult enterocytes, LREs rely on intracellular digestion within lysosomes to process macromolecular nutrients. This process demands a highly active endolysosomal system that breaks down complex proteins into absorbable units, which confers a particular architecture to LREs (**Figures S1A and S1B**)^9^. Autophagy, the cellular mechanism responsible for degrading and recycling cellular components, plays a central role in maintaining lysosomal function and turnover^25^. In LREs, elevated basal autophagy may support endolysosomal activity. Exploring this hypothesis could clarify the role of autophagy in the absorption and processing of nutrients in neonates, as well as its contribution to early development.

Utilising the GFP-LC3 transgenic mouse model (*GFP-Map1lc3a^TG/+^*) ^26^, we visualised autophagosomes as GFP-LC3 positive puncta and monitored autophagy flux via western blot (WB) by assessing the levels of free GFP, a byproduct of lysosomal degradation of GFP-LC3 II (**Figures 1A, 1B and S1C)**^27,28^. The presence and activity of LREs was determined by probing with mCherry/ fluorescent dextrans that are internalised as a macromolecule by LREs, but not by PECs **Figures 1A, 1B and S1C and S1D)** ^7^. Following gavage, fluorescent-dextran was rapidly internalised from the lumen into early and late endosomes (**Figure S1D**, arrowheads), accumulating progressively in lysosomal vacuoles (LVs) (**Figure S1D,** arrows). WB analysis was performed on lysates in epithelial cells from both proximal and distal sections of small intestines in GFP-LC3 mice at neonatal stages (P1, P7 and P14), and post-weaning (P21–30). This analysis revealed a proximal-distal gradient of protein absorption and autophagic degradation, mirroring the anterior-posterior intestinal maturation (**Figure 1B**)^29^.

**Figure 1.**
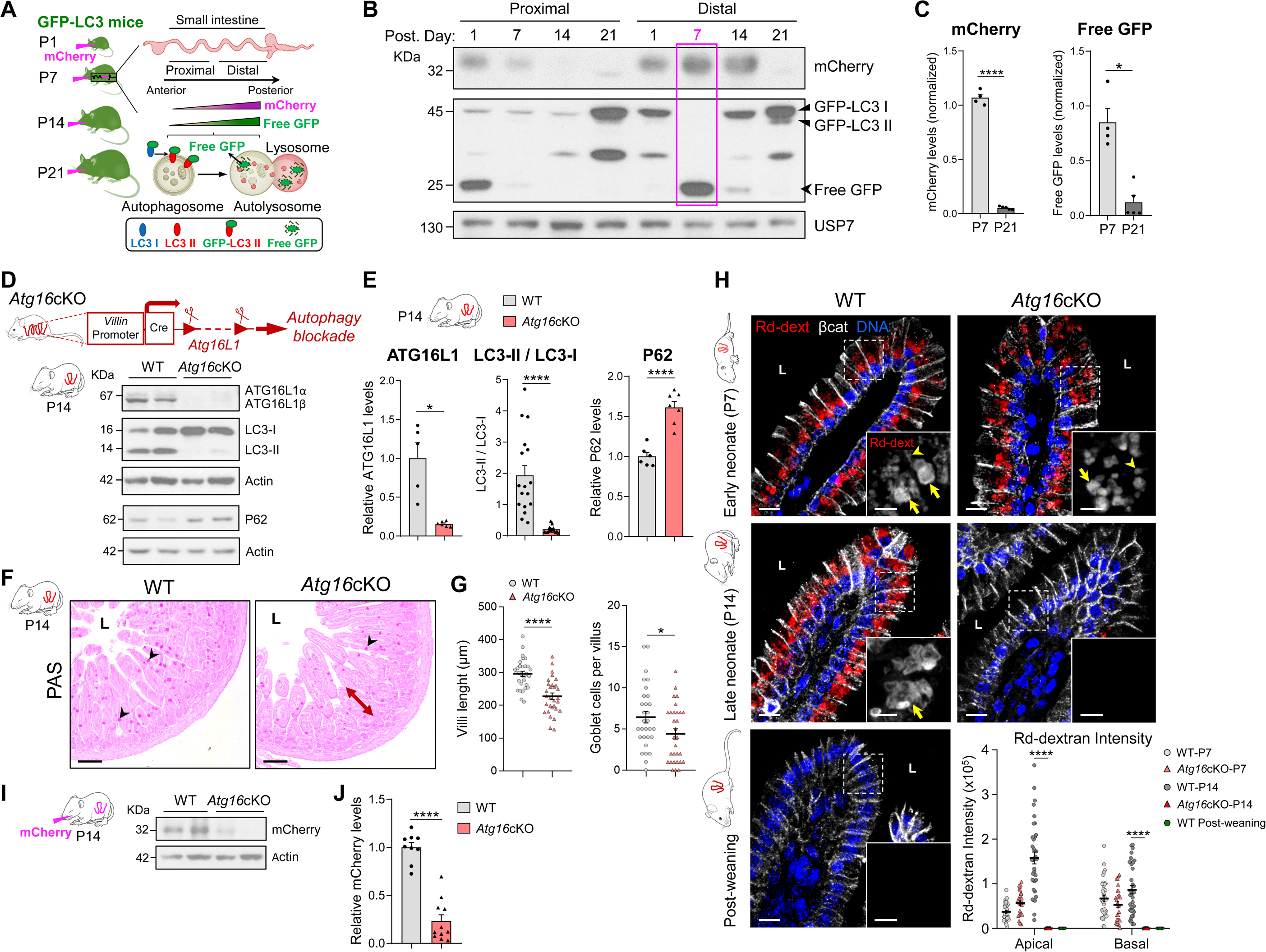
High Autophagic Activity of LREs Is Required for Nutrient Absorption. **(A and B)** Assessment of autophagic flux and protein absorption along the small intestine of GFP-LC3 mice during postnatal development. GFP-LC3 mice at postnatal days (P) 1, 7, 14, and 21 were orally gavaged with mCherry and sacrificed 3 hours later (n = 3) (A). Proximal and distal intestinal segments were harvested (A), and epithelial cell lysates analysed by immunoblotting for mCherry and GFP-LC3 isoforms (B). The representative western blot in (B) distinguishes cytosolic LC3-I, membrane-bound LC3-II, and cleaved “free GFP,” a marker of autolysosomal degradation. The pink rectangle highlights mCherry and free GFP in distal-P7, indicating maximal protein uptake and autophagic degradation. USP7, loading control. Figure 1A includes images adapted from Birchenough et al.^73^ (CC BY 4.0) and Zhao et al.^74^, and the GFP-LC3 isoforms scheme is based on Ni et al.^28^. **(C)** Quantification of mCherry and free GFP levels in lysates of ileal epithelial cells from P7 (n = 4) and P21 (n = 5) GFP-LC3 mice. Data derived from Figure S1C. **(D and E)** Generation and validation of *Atg16l1* intestinal knockout (*Atg16*cKO) mice. Ileal epithelial cell lysates from WT and *Atg16*cKO P14 mice were immunoblotted for ATG16L1, LC3, and P62 (D). (E) Quantification of ATG16L1 (n = 5,6) and P62 (n = 6,7), and LC3-II/LC3-I ratio (n = 17). In ATG16L1 and P62 graphs, data relative to WT. **(F and G)** Ileal architecture in *Atg16*cKO neonates. Periodic acid-Schiff (PAS)-haematoxylin staining of P14 ileal sections from WT and *Atg16*cKO mice (F) shows villus shortening (double red arrow) and a decrease in goblet cells (PAS+, arrowheads) in *Atg16*cKO sections (n = 3). L, lumen; scale bar, 100 µm. (G) Quantification of villus length and goblet cells per villus. Dots represent individual villi (n = 30 villi from 3 mice; 10 villi per mouse). **(H)** Confocal analysis of Rd-dextran uptake in ileal *Atg16*cKO enterocytes. WT and *Atg16*cKO neonates at P7 (early) and P14 (late), and post-weaning (P21–30) WT mice were gavaged with rhodamine B–dextran (Rd-dext, red) 16 hours prior to sacrifice (n = 3). Ileal sections labelled with β-catenin (βcat, white) and DAPI for DNA (blue). Dashed-line squares indicate magnified areas. Arrows point to lysosomal vacuoles; arrowheads, apical endosomes. L, lumen. Scale bars: 10 μm (main); 5 μm (insets). Dot plot shows the quantification of Rd-dextran intensity (integrated density) in the apical and basal regions of enterocytes. Dots represent individual enterocytes (n = 25–40). **(I and J)** *Atg16*cKO mice show impaired mCherry absorption. P14 mice were gavaged with mCherry and ileal epithelial cell lysates collected 16 hours later for immunoblotting (I). Actin, loading control. (J) Quantification of mCherry signal, normalised to WT (n = 9,11). In (C, E, J), dots represent individual mice and charts depict mean + SEM. In (G, H), dot plots show mean ± SEM (black lines). (C, E, G): Student’s t-test (mCherry, ATG16L1, P62, villus length) and Mann–Whitney test (free GFP, LC3 ratio, goblet cells/villus). (J): Mann–Whitney test. (H): Kruskal–Wallis test; only statistically significant comparisons between WT and *Atg16*cKO are shown within each region (apical or basal). *p < 0.05; ****p < 0.0001. **See also Figure S1.**

In the distal intestine, protein absorption preceded the induction of autophagy, with a subtle onset already detectable at P1, although no free GFP signal was observed. By P7, both mCherry uptake and free GFP increased markedly, with a strong accumulation of free GFP (**Figure 1B**, pink rectangle), indicating peak autophagic activity in LREs—coinciding with the establishment of neonatal intestinal features^30^. At P14, mCherry absorption persisted in the distal region, while free GFP levels declined, and by weaning (P21) both signals were absent (**Figure 1B)**. Quantification of mCherry and free GFP in ileal epithelial lysates at P7 and P21 confirmed these developmental differences **Figures 1C and S1C)**.

In contrast, the proximal intestine displayed a different temporal profile. At P1, both mCherry and free GFP were already present, but their levels progressively declined, and at P14 only low residual signals were detectable. Note that undegraded GFP-LC3 forms (I and II) and intermediate degradation products accumulated at P21, in both distal and proximal samples (**Figure 1B**).

Given these observations, we conclude that LREs maintain elevated autophagy levels relative to PECs, most probably as a result of their enhanced lysosomal function. To explore the consequences of autophagy disruption in these cells during early development, we utilised a genetic mouse model. ATG16L1 is crucial for both autophagosome formation and LC3 lipidation, two processes essential for autolysosomal degradation^31,32^. This gene has been extensively studied for its association with inflammatory bowel diseases^6,17^; nonetheless, its specific role in LRE functionality during the neonatal period remains largely unexplored. For this, we crossed with the *Atg16l1^fl/fl^*mice^33^, with mice expressing the *Villin-creER^T^*^2^ transgene^34^. This breeding strategy generated an *Atg16l1* knockout offspring in which the *Atg16l1* gene is conditionally deleted in the intestinal epithelium upon administration of tamoxifen (*Atg16*cKO) (**Figure 1D**).

Initially, we validated our model via WB analysis by measuring the levels of ATG16L1 and LC3 isoforms (cytoplasmic LC3-I and lipidated LC3-II) in the ileum of both WT and *Atg16*cKO mice at P14. We observed a significant reduction in ATG16L1 protein levels across all *Atg16*cKO samples, which led to a corresponding decrease in LC3-II/LC3-I ratio (**Figures 1D and 1E**). Specifically, our data showed an 84.5% reduction in ATG16L1 protein levels and an approximately 8-fold decrease in LC3-II/LC3-I ratio (**Figure 1E**). Furthermore, we observed elevated levels of P62, indicating a reduction in its autophagy-dependent degradation (**Figures 1D and 1E**). These outcomes are consistent with previous results^35^.

Subsequently, we examined the developmental trajectory of *Atg16*cKO mice, with a primary focus on their intestinal morphology and function. We observed that the mice reached the post-weaning stage (P30) without apparent issues, showing no weight defects throughout early postnatal development (**Figure S2A**). No significant changes were noted in the length or histological state of the small and large intestines (**Figures S2B-S2D**). However, a reduction in villus length and a decrease in the number of Goblet cells (GCs) were detected in the ileum at P14 (**Figures 1F and 1G**), along with mild oedema in the ileum at P30, an indicator of inflammation (**Figure S2D**). Inspired by these observations, we quantified specific intestinal epithelial (IEC) and immune cell subpopulations in the ileum of *Atg16*cKO mice at both neonatal and post-weaning stages using flow cytometry (**Figures S2E and S2F**). For this, we applied specific markers to identify various IEC subtypes, including enterocytes and their progenitors (EC), Goblet cells (GC), Paneth cells (PC), and enteroendocrine cells (EEC)^36^ (**Figure S3**, middle row). We also focused on cells related to innate immunity (IIC), such as CD11b+ cells (including NK cells, granulocytes, and macrophages/monocytes), as well as adaptive immunity, encompassing B lymphocytes (B cells) and CD4+ T helper cells (Th cells). We observed an increase in IIC and IEC populations during the neonatal period, accompanied by a concurrent decrease in GC and EEC in adult IECs (**Figure S2E**). In contrast, post-weaning samples showed a rise in B cells, stabilization in IIC and IEC levels, an increase in GC, and a decrease in PC and EEC (**Figure S2F**). These results corroborate our histological findings regarding GCs and inflammation.

Next, we investigated the effect of autophagy inhibition on the absorption of macromolecules in the ileum. We gaveged rhodamine B–dextran (Rd-dextran) as a probe in *Atg16*cKO and WT mice as controls (**Figure 1H**). Our analysis differentiated between the apical and basal regions of enterocytes to assess the sequential entry of Rd-dextran, according to the internalization dynamics observed in these cells(**Figure S1D**). Confocal microscopy demonstrated that at P7, *Atg16*cKO enterocytes internalized Rd-dextran similarly to WT counterparts. Nevertheless, it is worth noting that in *Atg16*cKO enterocytes, Rd-dextran levels were slightly elevated in the apical region but reduced in the basal region compared to WT (**Figure 1H**). This observation suggests a deceleration in the internalisation of macromolecules during this developmental period. Subsequently, at P14, *Atg16*cKO enterocytes exhibited a complete abrogation of macromolecule absorption, resembling PECs and indicating an advancement in intestinal maturation (**Figure 1H**). To validate the observed decrease in absorption at P14, we probed neonates from both WT and *Atg16*KO groups with mCherry and quantified the internalised mCherry in the ileum via WB. The results confirmed consistently lower levels of nutrient absorption in *Atg16*KO mice (**Figures 1I and 1J**). Altogether, we conclude that blocking autophagy severely disrupts macromolecule absorption in the neonatal ileum.

To further investigate this role autophagy in macromolecule absorption, we examined the expression of the endocytic genes *Dab2*, *Amn*, and *Mamdc4*, as well as the lysosomal genes *Mcoln3*, *Lamp2*, and *Ctsl1*. Previous studies have demostrated the specific role of these genes on the endolysosomal function of LREs^7,12,13^. Indeed, we validated their expression during early postnatal intestinal development (**Figure 2A**). As expected, an upregulation of these genes was observed in distal samples at P7, coinciding with the presence of LREs (**Figure 2A**). Moreover, a proximodistal expression pattern was observed, consistent with the antero-posterior maturation of the intestine^29^ (**Figure 2A**).

**Figure 2.**
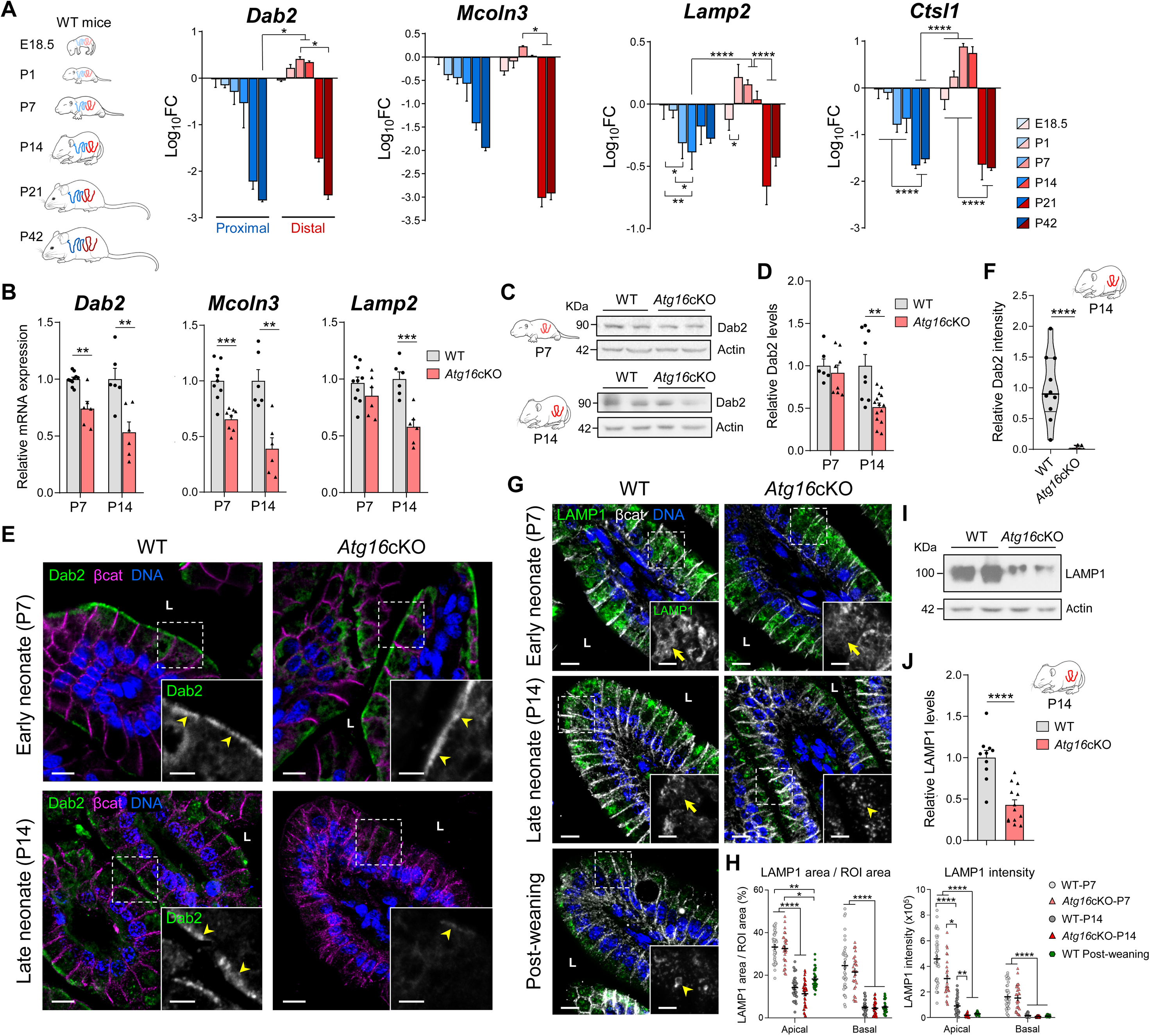
Loss of Autophagy Compromises the Endolysosomal System in LREs. **(A)** Reverse transcription quantitative polymerase chain reaction (RT-qPCR) analysis of *Dab2*, *Mcoln3*, *Lamp2* and *Ctsl1* genes in epithelial cells from proximal and distal samples of small intestines from WT mice at the indicated time points (n = 3). Data relative to values of proximal region of small gut from embryonic day (E) 18.5 mice embryos, and presented as log_10_ fold changes (log_10_FC). **(B)** RT-qPCR of *Dab2*, *Mcoln3*, and *Lamp2* in ileal epithelial cells from WT and *Atg16*cKO mice at P7 and P14. Sample sizes: *Dab2* and *Lamp2* (n = 10,6,6,6), *Mcoln3* (n = 9,8,6,6). **(C and D)** Immunoblot analysis of Dab2 protein in ileal epithelial cell lysates from WT and *Atg16*cKO mice at P7 and P14 (C). Actin, loading control. (D) Quantification of Dab2 protein levels (n = 6,8,9,14). **(E and F)** Confocal images of ileal sections from WT and *Atg16*cKO mice at P7 and P14 (n = 3) labelled with anti-Dab2 (green), β-catenin (magenta), and DAPI (blue) (E). Arrowheads mark apical Dab2 localization. (F) Violin plot quantifying apical Dab2 intensity (integrated density) of P14 sections, normalised to epithelial area (n = 10,8; 2–4 images per mouse). **(G and H)** Lysosomal occupation and content in WT and *Atg16*cKO mice at P7 and P14, and in post-weaning (P21–30) WT mice. Confocal sections labelled with anti-LAMP1 (green), β-catenin (white), and DAPI (blue) (G). Arrows: endolysosomal system in WT LREs; arrowheads: reduced LAMP1^+^ vesicles in *Atg16*cKO and post-weaning enterocytes. (H) Dot plots quantify the percentage of LAMP1^+^ area and LAMP1 intensity (integrated density) within apical and basal regions of interest (ROIs) (n = 26–36). **(I and J)** Immunoblot analysis of LAMP1 protein in ileal epithelial cell lysates from WT and *Atg16*cKO mice at P14 (I). Actin, loading control. (J) Quantification of LAMP1 protein levels (n = 10,13). In (E, F), dashed-line squares indicate magnified areas; L, lumen. Scale bars: 10 μm (main); 5 μm (insets). Dots represent individual mice (B, D, J) or images (F) or cells (H). In (B, D, F, J), data relative to WT values. Graphs of (A) represent mean + SD; graphs of (B, D, J), mean + SEM; violin plot of (F), median (thick line) and interquartile range (IQR, thin lines); dot plots of (H), mean ± SEM (black lines). Statistical analyses: Kruskal–Wallis test (A [*Dab2*, *Mcoln3*], H), two-way ANOVA (A: *Lamp2*, *Ctsl1*), Student’s *t*-test (B, D, J), Mann–Whitney test (F). In (A), only the most relevant statistically significant comparisons are shown. In (H), only statistically significant comparisons are displayed within each group (apical or basal). *p < 0.05, **p < 0.01, ***p < 0.001, ****p < 0.0001. **See also Figure S2.**

Then, we studied the mRNA expression of *Dab2*, *Mcoln3*, and *Lamp2* genes in both the proximal and distal regions of the small intestines of WT and *Atg16*cKO mice at P7 and P14 (**Figures 2B and S2G**). Since LREs predominantly localise in the distal region, the proximal segment was also analysed to assess potential compensatory mechanisms. In distal *Atg16*cKO samples at P7, mRNA levels of these LRE markers were significantly reduced (**Figure 2B**), while the proximal region showed no significant changes compared to WT mice (**Figure S2G**). By P14, a substantial reduction in gene expression was evident in both proximal and distal *Atg16*cKO mice samples (**Figures 2B and S2G**). Additionally, we analysed *Amn*, *Mamdc4,* and *Ctsl1* mRNA expression in the ileum at P14, finding lower expression levels in the *Atg16*cKO samples (**Figure S2H**).

To delve deeper into the protein levels of Dab2, we performed WB and immunofluorescence (IF) analysis in the ileum of *Atg16*cKO mice at P7 and P14. While P7 samples exhibited reduced mRNA expression (**Figure 2B**), Dab2 protein levels remained comparable to WT; however, by P14, a marked reduction in Dab2 was observed in *Atg16*cKO samples (**Figures 2C-2F**).

Furthermore, we assess the lysosomal occupancy and content in the apical and basal regions of WT and *Atg16*cKO ileal enterocytes using confocal microscopy with an anti-LAMP1 antibody as a lysosomal marker, as the *Lamp1* and *Lamp2* genes exhibit similar expression patterns (data not shown). Post-weaning (P21– 30) WT mice were used as controls. At P7, both WT and *Atg16*cKO enterocytes exhibited comparable lysosomal occupancy and content, with peak intensity observed in both conditions (**Figures 2G and 2H**). However, by P14, while lysosomal occupancy remained similar, *Atg16*cKO enterocytes demonstrated reduced intensity in both the apical and basal regions (**Figures 2G and 2H**). Magnified images also revealed that lysosomes in *Atg16*cKO enterocytes were smaller in P14 neonates, resembling those of PECs (**Figure 2G**, arrowheads). This reduction in lysosomal content in *Atg16*cKO P14 ileum was further confirmed by WB analysis (**Figures 2I and 2J**).

In summary, our results indicate that autophagy plays a crucial role in preserving endolysosomal homeostasis of LREs. Disruption of autophagy in LREs using the *Atg16*cKO mice results in significant defects in macromolecule uptake and lysosomal function, accompanied by the downregulation of key endocytic and lysosomal genes, particularly in the distal ileum of late neonatal mice. These outcomes highlight the importance of autophagy in maintaining LRE function and, consequently, ensuring proper postnatal development. However, the precise mechanisms by which autophagy exerts its effects remain unclear.

### LREs show a significant induction of mitophagy to control mitochondrial homeostasis

To elucidate the primary type of autophagy in neonatal enterocytes, we defined the gene expression signature of LREs through bulk RNA sequencing. Leveraging the high levels of macromolecular absorption and autophagy in LREs, we probed neonatal GFP-LC3 mice (P7) with Rd-dextran and isolated LREs from the cellular suspension of the ileum by fluorescence-activated cell sorting (FACS), selecting them as the epithelial cells with high levels of autophagy and Rd-dextran (Rd-dext^+^GFP^high^). Moreover, to highlight their genetic neonatal signature, LREs were compared to post-weaning enterocytes (PECs) from mice (P42). Following RNA extraction, we conducted next-generation sequencing (NGS) analysis (**Figures 3A and S4A**).

**Figure 3.**
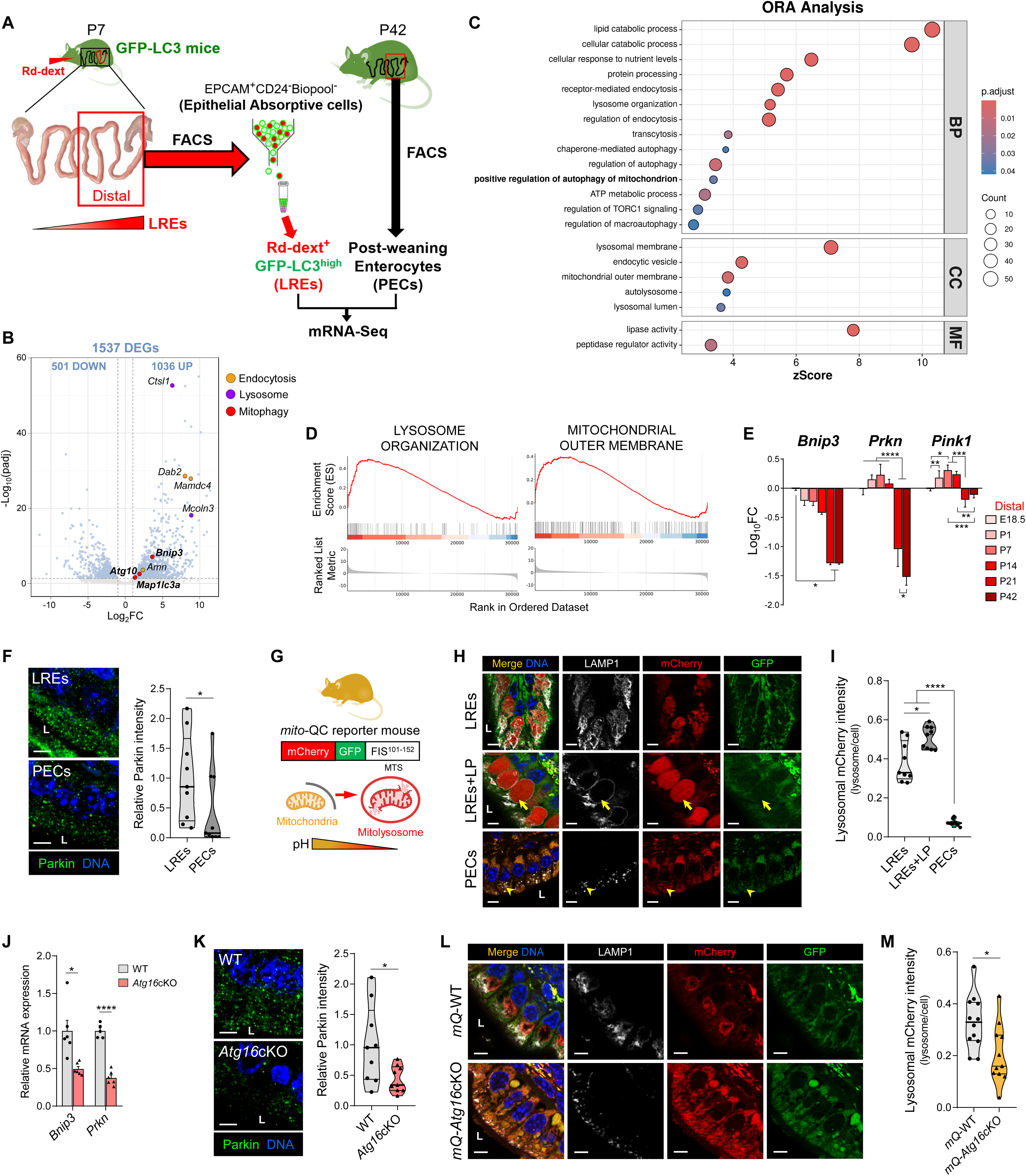
LREs Present High Levels of Mitophagy. **(A)** Workflow of RNA sequencing (RNA-seq) analysis comparing LREs from P7 GFP-LC3 mice gavaged with rhodamine B–dextran (Rd-dext) (16 hours prior) to PECs from P42 GFP-LC3 mice. Enterocytes were isolated from ileal samples by fluorescence-activated cell sorting (FACS) and used for RNA-seq. See Figure S4A for gating strategy. Scheme includes images adapted from Treuting et al.^75^ and Park et al.^7^. **(B)** Volcano plot of differentially expressed genes (DEGs) between LREs and PECs. Each point represents a gene, plotted by log2 fold change (x-axis) and –log10 adjusted p-value (padj) (y-axis). DEGs (n = 1537; shown in steel blue) were defined as genes with padj < 0.05 and |log2FC| > 1. Upregulated genes associated with endocytosis (amber), lysosome (purple), and mitophagy (red) are highlighted. Dashed lines indicate significance (padj < 0.05) and fold-change thresholds (±1). Genes with –log10(padj) > 60 are not shown. **(C)** Bubble plot of selected Gene Ontology (GO) terms from over-representation analysis (ORA) of the differential gene expression data from the LREs vs PECs. Note the presence of regulation of autophagy, positive regulation of autophagy of mitochondrion and mitochondrial outer membrane pathways. The x-axis represents the z-score, bubble size reflects the number of associated genes, and bubble colour indicates statistical significance (adjusted p-value), with a red-to-blue gradient from most to least significant. GO terms are grouped into three categories: BP, biological process; CC, cellular component; MF, molecular function. **(D)** Enrichment plots of lysosome organization and mitochondrial outer membrane gene sets from gene set enrichment analysis (GSEA) comparing LREs and PECs. The top panel shows the Enrichment Score (ES) across ranked genes, with the peak indicating maximum enrichment. The middle panel marks gene positions in the ranked list. The bottom colour bar reflects gene correlation with phenotype (red: positive, blue: negative). **(E)** RT-qPCR analysis of *Bnip3*, *Prkn*, and *Pink1* genes in ileal epithelial cells from WT mice at indicated time points (n = 3). Values are expressed as log10 fold change relative to E18.5 embryonic levels. **(F)** Confocal images of ileal sections from WT mice at P7 (LREs) and P21–30 (PECs), labelled with anti-Parkin (green) and DAPI (blue). Left: violin plot of Parkin intensity (integrated density) normalised to epithelial area. **(G)** Schematic of the *mito*-QC reporter mouse model. Mitochondria are tagged with the chimeric protein mCherry-GFP-FIS1^101–152^. Upon mitophagy, mitochondria are delivered to the lysosome where acidic pH quenches GFP but not mCherry, allowing identification of mitolysosomes as mCherry-only puncta. **(H)** Confocal images of ileal sections from *mito*-QC mice at P7 (LREs) (± leupeptin, LP) and P21–30 (PECs). Samples stained with anti-LAMP1 (white) and DAPI (blue). Arrows indicate enlarged, blocked lysosomes with red-only mitochondrial aggregates in LREs+LP; arrowheads point to LAMP1⁺/mCherry⁺ puncta lacking GFP, validating the *mito*-QC model. **(I)** Violin plot of lysosomal mCherry intensity normalised to cellular mCherry intensity from (H), based on integrated density measurements. **(J)** RT-qPCR of *Bnip3* and *Prkn* in ileal epithelial cells from WT and *Atg16*cKO mice at P14. Dots represent individual mice. Sample sizes: *Bnip3* (n = 6, 6), *Prkn* (n = 5, 6). **(K)** Confocal images of ileal sections from WT and *Atg16*cKO mice at P14 stained with anti-Parkin (green) and DAPI. Left: violin plot of Parkin intensity (Integrated Density) normalised to epithelial area. **(L)** Confocal images of ileal sections from *mQ-*WT and *mQ-Atg16*cKO mice at P14 (n = 3), labelled with anti-LAMP1 (white) and DAPI (blue). **(M)** Violin plot of lysosomal mCherry intensity normalised to cellular mCherry intensity from (L), based on integrated density measurements (n = 12–11; 3–4 images per mouse). In images of (F, H, K, L), L, lumen. Scale bars: 5 μm. Images of (H, L) deconvolved with Huygens software. In (A–I) and (K–M), n = 3; in (F, I, K), n = 9 images from 3 mice (3 images per mouse). In graphs of (F, I), data relative to LREs values; in (J, K, M), relative to WT values. In (E), graph represents mean + SD; in (J), mean + SEM. In violin plots of (F, I, K, M), the median (thick line) and IQR (thin lines) are shown and dots represent individual images. Statistical analyses: Kruskal–Wallis test (E: *Bnip3*), one-way ANOVA (E [*Prkn*, *Pink1*], I), Student’s *t*-test (F, J, K, M). *p< 0.05, **p< 0.01, ***p < 0.001. **See also Figure S4.**

We detected a total of 1,537 differentially expressed genes (DEGs), of which 1,036 were upregulated and 501 were downregulated in LREs (**Figure 3B**). Differential expression analysis and principal component analysis demonstrated that each population exhibited unique enrichment signatures (**Figures S4B and S4C**). The computational method of over-representation analysis (ORA) identified which biological processes were overrepresented in the differentially expressed genes (DEGs)^37^. The majority of differentially regulated pathways were associated with lysosomal function, nutrient absorption, processing, and transport (**Figures 3C, 3D, and S4D**). These findings are consistent with earlier RNA-Seq data from LREs in zebrafish^7^. Nonetheless, the regulation of lipid metabolism pathways differed (**Figure 3C and S4D**), suggesting that mammalian LREs significantly modulate these pathways to process lipids from maternal milk.

Among the most upregulated genes were those associated with endocytosis, such as *Mamdc4*, *Dab2*, and *Amn*, as well as lysosome-associated genes like *Mcoln3* and *Ctsl1* (**Figure 3B**). The upregulation of *Dab2*, *Amn*, and *Lamp2* genes corresponded with previously published ileum transcriptome data from mice and zebrafish^7^. Additionally, the overexpression of *Mamdc4* and *Mcoln3* genes also aligned with earlier results in LREs from rodents^12,38^. We confirmed elevated levels of *Ctsl1* mRNA expression and lysotracker staining in LREs compared to PECs in WT mice (**Figures S4E and S4F**), which validated the RNA-Seq data and ensured that the lysosome-associated pathway was not positively regulated due to the selection of neonatal enterocytes with high autophagy levels.

Interestingly, the ’positive regulation of autophagy of mitochondrion ’ and ’mitochondrial outer membrane’ pathways appeared significantly enriched in LREs (**Figures 3C and 3D**). These outcomes, along with the upregulation of several genes involved in mitophagy, such as *Bnip3*, *Atg10*, and *Map1lc3a* (**Figure 3B**)^39,40^, indicated that mitophagy is a significant component of the elevated autophagic activity observed in LREs. To explore this hypothesis, we examined the mRNA expression of the canonical mitophagy genes *Prkn* (Parkin) and *Pink1*^41^, together with *Bnip3* as a control, in ileal samples from WT mice across developmental stages (E18.5-P42). While *Bnip3* exhibited a progressive decrease in expression throughout development, *Prkn* and *Pink1* showed upregulation in neonatal samples, especially *Prkn* (**Figure 3E**). Moreover, we observed that LREs displayed increased Parkin protein levels compared to PECs through IF (**Figure 3F**).

Next, we assessed mitophagy in LREs and PECs using the fixable *mito*-QC reporter C57BL/6J mouse^42^, which ubiquitously expresses a chimeric protein consisting of pH-insensitive mCherry, a pH-sensitive GFP, and the mitochondria-targeting sequence (MTS) from the outer mitochondrial membrane protein FIS1 (amino acids 101–152). This allowed us to differentiate between cytoplasmic mitochondria (mCherry^+^GFP^+^) and mitochondria degraded by mitophagy in lysosomes (mCherry^+^GFP^−^) (**Figure 3G**). Confocal images of LREs revealed a high concentration of red-fluorescent mitochondria inside their endolysosomal system (**Figures 3H and 3I**). In fact, when we inhibited autophagic flux by treating the neonates with the protease inhibitor Leupeptin (40 mg/kg for 16 hours)^43^, we observed a significant expansion of LVs and a considerable accumulation of red-only mitochondria within them (**Figure 3H** arrows; **and 3I**). In contrast, PECs exhibited fewer red mitochondria in lysosomes and a uniform distribution of yellow mitochondria throughout the cytoplasm (**Figures 3H and 3I**).

Thus, we confirmed the high mitophagic activity of LREs, suggesting severe impairment of mitophagy and mitochondrial integrity in *Atg16*cKO enterocytes. Indeed, we found that mRNA expression levels of both *Bnip3* and *Prkn* (**Figure 3J**), along with Parkin protein levels (**Figure 3K**), were significantly lowered in neonatal *Atg16*cKO enterocytes. To further investigate the robust mitophagic activity of LREs, we generated an autophagy-deficient *mito*-QC mouse line (*mQ-Atg16*cKO). We observed that in *Atg16*cKO LREs exhibited reduced mitochondrial mass within the lysosomes (**Figures 3L and 3M**), consistent with the low mitophagy observed in PECs (**Figures 3H**, lower panels). Therefore, the suppression of autophagy in LREs results in the downregulation of mitophagy-related genes and a reduced uptake of mitochondria into lysosomes.

Motivated by these results, we addressed the mitochondrial status of *Atg16*cKO mice. Using transmission electron microscopy (TEM), we analysed mitochondrial damage, quantity, and morphology in ileal enterocytes from neonatal *Atg16*cKO mice (**Figure 4A-4C**). Mitochondria were categorized into three classes based on electron density, cristae appearance, matrix loss, and outer membrane integrity: class I (normal), class II (partially affected), and class III (damaged) (**Figure 4B**)^44^. For each class, the number, percentage, area, and dimensional ratio (aspect ratio, AR: major axis/minor axis) were quantified (**Figure 4C**). The data showed that number and percentage of mitochondria within each class were similar across both WT and *Atg16*cKO enterocytes. However, while class I mitochondria in *Atg16*cKO enterocytes exhibited a smaller area and a higher AR, class III mitochondria showed a larger area, suggesting an accumulation of damaged mitochondrial mass due to impaired autophagy (**Figure 4C**).

**Figure 4.**
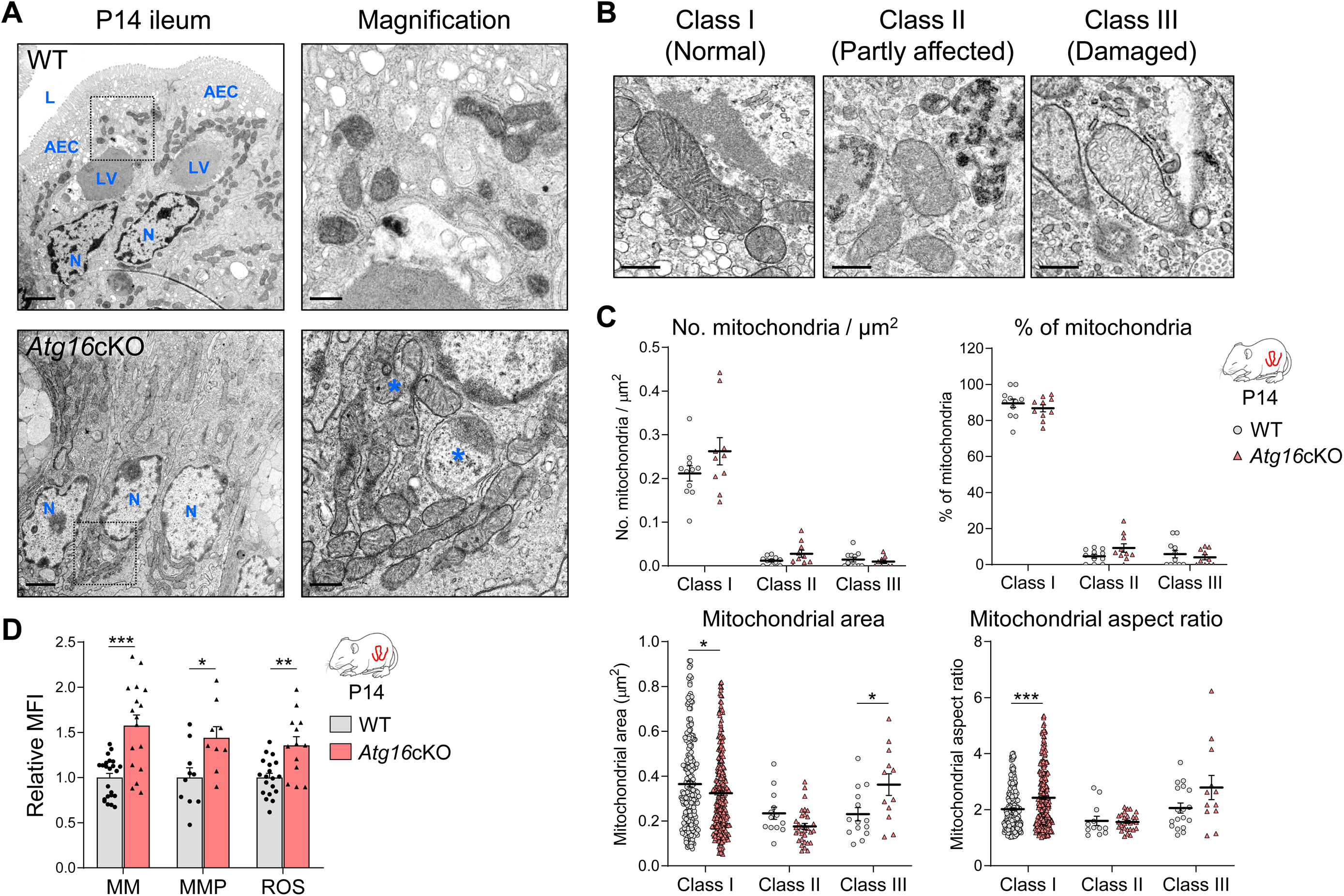
*Atg16*cKO Neonatal Mice Exhibit Increased Mitochondrial Damage and Oxidative Stress in Enterocytes. **(A)** Representative transmission electron microscopy (TEM) micrographs of ileal sections from WT and *Atg16*cKO mice at P14 (n = 3). Note the reduction of the endolysosomal system and the absence of lysosomal vacuole in the *Atg16*cKO enterocytes. Dashed boxes indicate magnified areas. N, nucleus; LV, lysosomal vacuole; AEC, apical endosomal complex; L, lumen. Asterisks indicate damaged mitochondria. Scale bars: 2 μm (main); 500 nm (insets). **(B)** TEM-based classification of mitochondria into three categories based on electron density and cristae integrity: Class I (normal), Class II (partially affected), and Class III (damaged). Classification based on Moschandrea et al. ^44^. See Electron Microscopy Section in Methods. Scale bar: 500 nm. **(C)** Distribution of mitochondrial integrity classes in *Atg16*cKO neonatal enterocytes. Quantification of the number, percentage, area and aspect ratio (long axis / short axis) of each mitochondria class in (A), as defined in (B). In count and percentage graphs, dots represent individual enterocytes; in area and aspect ratio graphs, dots represent individual mitochondria. Sample sizes are indicated in Table S1. **(D)** Flow cytometry analysis of mitochondrial status of *Atg16*cKO neonatal enterocytes. Graphs show relative mean fluorescence intensity (MFI) for Mitotracker Green (mitochondrial mass, MM), Mitotracker Red CMXROS (mitochondrial membrane potential, MMP) and CM-H_2_DCFDA (ROS) in ileal EpCAM^+^CD24^-^ cKit^-^ cells from WT and *Atg16*cKO mice at P14. See Figure S3 for gating strategy. Dots represent individual mice (MM, n = 23,17; MMP, n = 10,9; ROS, n = 19,13). In (C), dot plots represent mean ± SEM (black lines). In (D), bar chart show mean + SEM. Statistics analyses: Mann–Whitney test (C, D [MM]), Student’s *t-*test (D: MMP, ROS). *p < 0.05, **p < 0.01, ***p < 0.001, ****p < 0.0001. **See also Figure S3 and Table S1.**

To further validate the mitochondrial status, we performed flow cytometry on neonatal *Atg16*cKO enterocytes using mitochondrial probes *ex vivo*: Mitotracker Green for mitochondrial mass, Mitotracker Red CMXROS for membrane potential, and CM-H_2_DCFDA for ROS^36,45^. This analysis confirmed an increase in mitochondrial mass, membrane potential, and ROS in *Atg16*cKO enterocytes (**Figure 4D**). In conclusion, the inhibition of mitophagy in LREs leads to an increased in the mass of dysfunctional mitochondrial.

Collectively, mitophagy represents a significant component of the heightened autophagic activity observed in LREs. Its inhibition not only contributes to the mitochondrial defects seen in neonatal *Atg16*cKO enterocytes but may also account for the impairment of their endolysosomal system. These findings highlight a compelling link between the mitochondria-lysosomes crosstalk and neonatal development, as well as the associated metabolic adaptations.

### Dysfunction of the mitochondria-lysosome axis in the neonatal epithelium causes an early metabolic switch dependent on BLIMP1 and MAFB

An interesting phenotype observed in neonatal *Atg16*cKO enterocytes was their resemblance to PECs, particularly in terms of the abrogation of macromolecule absorption and the reduction in endolysosomal system functionality. This phenotype could suggest an acceleration of their metabolic transition. At the end of lactation, the mammalian intestine becomes capable of digesting nutrients extracellularly, replaces LREs with PECs (without LVs), and regulates the expression of enzymes to adapt to the new solid diet^3,46^. Previous studies have demonstrated that the transcriptional factors BLIMP1, MAFB, and c-MAF (two members of the Maf family of proteins) mediate the metabolic shift in enterocytes during neonatal development^47–49^. At weaning, Blimp1 expression decreases, leading to a downregulation of neonatal enzymes such as β-galactosidase (Glb1) and argininosuccinate synthase 1 (Ass1), and an increase in expression of adult enzymes, including sucrase-isomaltase (Sis), trehalase (Treh), and arginase 2 (Arg2)^48,49^. Similarly, early inhibition of MAFB and c-MAF in neonatal intestine induces a comparable metabolic switch ^47^.

Our RNA-Seq analysis comparing LREs and PECs confirmed these dynamics: LREs expressed higher levels of *Mafb*, *Blimp1*, *Glb1*, and *Ass1*, along with a decrease of *Sis*, *Treh*, and *Arg2* (**Figure S5A**). *c-maf* showed no significant differences (**Figure S5A**), though transient regulation at specific neonatal stages cannot be excluded. Consistently, RT-qPCR analysis of proximal and distal small intestine samples from WT mice across development (E18.5–P42) revealed neonatal enrichment of transcription factors and neonatal enzymes, whereas post-weaning samples displayed the opposite profile, with strong induction of adult enzymes (**Figures S5B and S5C**). *Arg2* expression levels closely paralleled those of Sis and Treh (data not shown).

To investigate whether autophagy deficiency contributes to this transition, we studied the expression of these metabolic genes in our *Atg16*cKO model. Gene expression analysis revealed a moderate downregulation of the transcription factor genes *Blimp1* and *Mafb*, except for *c-maf*, as well as a more pronounced downregulation of neonatal enzyme genes *Ass1* and *Glb1*. In contrast, there was an upregulation of adult enzyme genes *Treh*, *Sis*, and *Arg2* (**Figure 5A**). Similarly, confocal microscopy analysis revealed reduced levels of MAFB, BLIMP1 (**Figures 5B and 5C**), and ASS1, along with increased levels of TREH in neonatal *Atg16*cKO enterocytes (**Figures 5D and 5E**). This later result was also confirmed by WB analysis in neonates (**Figure SF and SG**). In summary, *Atg16*cKO mice advance their metabolic transition in enzymes regulated by MAFB/BLIMP1 in their neonatal epithelium.

**Figure 5.**
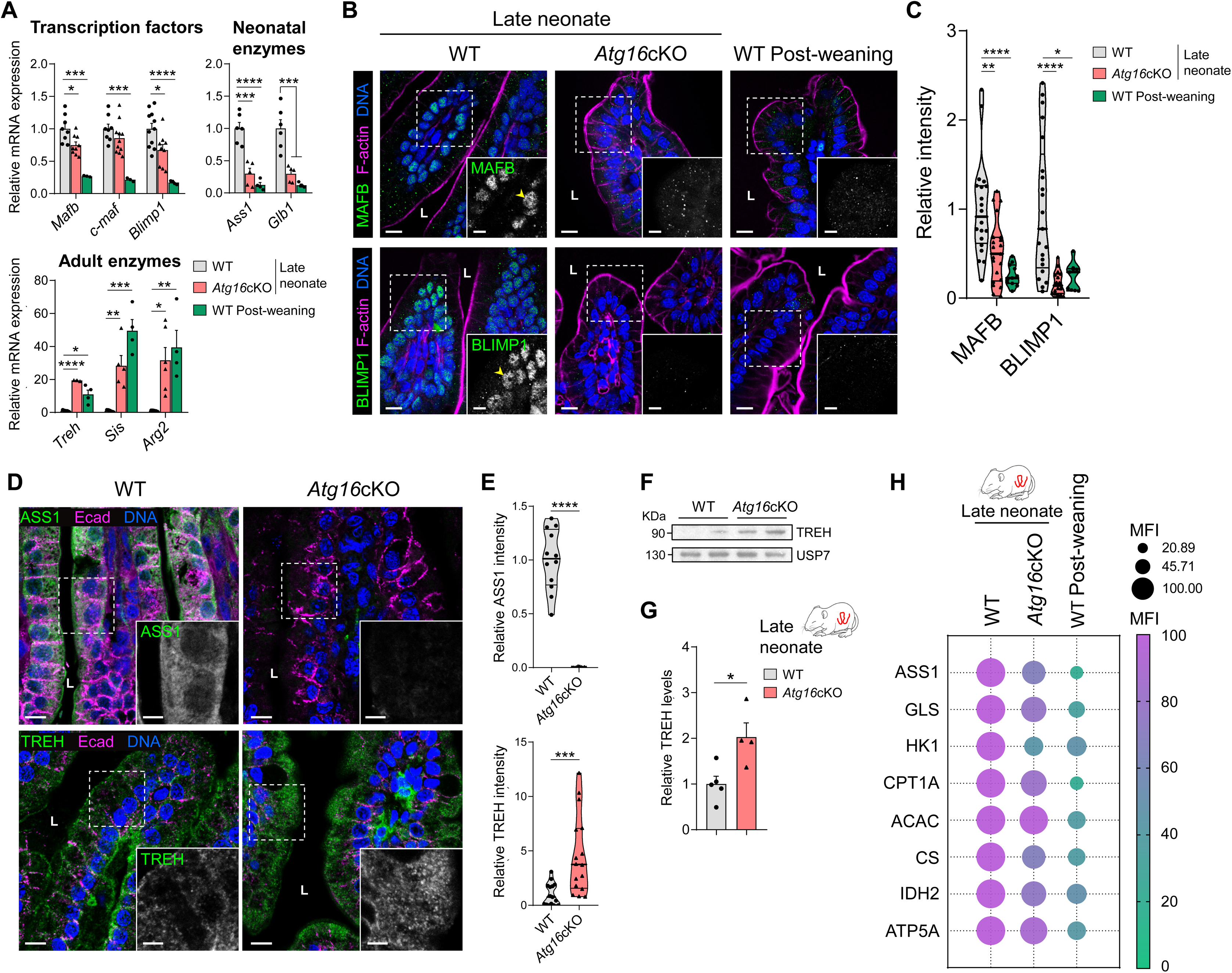
Autophagy Deficiency Causes Early Metabolic Transition in LREs. **(A)** RT-qPCR analysis of transcription factor (*Mafb*, *c-maf*, *Blimp1*), neonatal (*Ass1*, *Glb1*) and adult (*Treh*, *Sis*, *Arg2*) enzyme genes in ileal epithelial cells from WT and *Atg16*cKO mice at late neonatal stage (P14) and from WT mice at post-weaning stage (P21–30). Sample sizes: *Mafb* (n = 8, 9, 3), *c-maf* (n = 8, 11, 3), *Blimp1* (n = 11, 9, 4), neonatal enzymes (n = 6, 5, 4), *Treh* (n = 5, 3, 4), *Sis* (n = 5, 5, 4), *Arg2* (n = 5, 6, 4). **(B)** Confocal images of ileal sections from WT and *Atg16*cKO mice at P14 (n = 5), and from post-weaning (P21–30) WT mice (n = 2–3), labelled with anti-MAFB (green, top) or anti-BLIMP1 (green, bottom), and with phalloidin for F-actin (magenta) and DAPI (blue). Arrowheads point to nuclear localization of MAFB and BLIMP1 in WT LREs. **(C)** Violin plot of nuclear MAFB (n = 20, 21, 11) and BLIMP1 (n = 21, 16, 9) intensity (integrated density) in (B), normalised to epithelial nuclear area. **(D)** Confocal images of ileal sections from WT and *Atg16*cKO mice at P14, labelled with anti-ASS1 (green, top) (n = 3) or anti-TREH (green, bottom) (n = 5), anti-E-cadherin (Ecad, magenta), and DAPI (blue). **(E)** Violin plots of ASS1 (top; n = 12, 10) and TREH (bottom; n = 15,16) intensity (integrated density) in (D), normalised to epithelial area. **(F)** Western blot of TREH protein from ileal epithelial cells of WT and *Atg16*cKO mice at P14. USP7, loading control. **(G)** Bar chart quantifying TREH protein levels from (F) (n = 5, 4). **(H)** Met-Flow profiling of ileal EpCAM^+^CD24^-^ cells from WT (n = 10–13) and *Atg16*cKO (n = 7–9) mice at P14, and from post-weaning (P21–30) WT mice (n = 8). Bubble colour and size reflect group means for each marker. See Figure S3 for the gating strategy and Table S2 for statistical significance. In (B, D), dashed boxes indicate magnified areas; L, lumen. Scale bars: 10 μm (main); 5 μm (insets). In (A, C, E, G, H), data relative to WT P14 values. In (A, G), graphs show mean + SEM and dots represent individual mice. In (C, E), violin plots depict the median (thick line) and IQR (thin lines) and dots are individual images (2–5 images per mouse, 2–5 mice per group). Statistical analyses: one-way ANOVA (A, C [MAFB], H), Kruskal–Wallis test (C: BLIMP1), Mann–Whitney test (E), Student’s t test (G). *p < 0.05, **p < 0.01, ***p < 0.001, ****p < 0.0001. **See also Figures S3 and S5, and Table S2.**

Next, we studied the metabolic profiles of neonatal enterocytes from *Atg16*cKO mice, in comparison with neonatal and post-weaning WT mice. In order to evaluate deeply the metabolic transition during the neonatal development and the impact of autophagy, we used a pioneer technique called Met-flow. The Met-Flow technique is a flow cytometry-based method that examines the metabolic state of cells by evaluating the levels of rate-limiting enzymes across key metabolic pathways. Specifically, we studied ASS1 and GLS as markers of amino acid (AA) metabolism, HK1 as a marker of glycolysis, CPT1A and ACAC as markers of fatty acid (FA) metabolism, CS and IDH2 for the tricarboxylic acid (TCA) cycle, and ATP5A for oxidative phosphorylation (OXPHOS)^45,50^ (**Figure S5D**). Met-Flow analysis revealed higher expression of most metabolic pathways in LREs compared to PECs (**Figure S5E**), reflecting a their hyperconsumptive signature aligned with the high energy demands of this developmental period^51^. Notably, neonatal *Atg16*cKO enterocytes exhibited a moderate reduction in enzyme protein levels, positioning them between those of WT neonates and post-weaning mice (**Figure 5H**). This intermediate profile is indicative of a globally accelerated metabolic transition toward a more adult state.

In summary, our results reveal that autophagy is essential for coordinating mitochondrial and metabolic remodeling during neonatal ileum development and the weaning transition. Suppression of autophagy perturbs endolysosomal and mitochondrial integrity, thereby altering the transcriptional control of LREs. Consequently, LREs undergo a premature metabolic switch, marked by MAFB and BLIMP1 dysregulation and reduced activity of key metabolic pathways.

### Mitochondrial homeostasis is critical for maintaining LREs function and their metabolic transition

After identifying mitochondrial homeostasis defects as a downstream consequence of impaired endolysosomal function in the *Atg16*cKO model, we next investigated whether mitochondria themselves play an intrinsic role in sustaining LRE integrity. For this purpose, we used the *Tfam*cKO mouse model (*Tfam^fl/fl^/Villin-creER^T^*^2^), which enables inducible deletion of Tfam in the neonatal intestinal epithelium. This approach allowed us to directly test whether mitochondrial dysfunction is sufficient to compromise lysosomal homeostasis in LREs.

TFAM is a transcription factor that regulates the transcription and replication of mitochondrial DNA (mtDNA)^52^. While the role of mitochondria in maintaining intestinal homeostasis has been extensively explored^24^, and the consequences of Tfam deletion have been reported in both embryonic and adult intestinal epithelium^53^, its specific contribution during the neonatal stage, and particularly in nutrient absorption and processing by LREs, remains unexplored.

We first characterized the general phenotype of *Tfam*cKO pups during the neonatal period. These mice exhibited developmental defects, a generally poor appearance, and significantly reduced body weight, starting around P8, which ultimately compromised their survival and prevented successful weaning (**Figures 6A-6C**). Given the severity of the phenotype and in adherence with animal welfare guidelines, euthanasia was performed after two consecutive days without weight gain or upon observing weight loss of around 10% (**Figure 6C**). Based on the timing of euthanasia, we defined two experimental groups: mice euthanised between P9–P11 (hereafter referred to as P10), and those euthanised between P13–P15 (hereafter referred to as P14). In the ileum of both groups, we confirmed a marked reduction in *Tfam* mRNA expression (**Figure 6D**), accompanied by decreased levels of mtDNA, as assessed through the quantification of electron transport chain-encoded genes including *mt-Nd1*, *mt-Co*1, *mt-Co2*, and *mt-Atp6* (**Figure S6A**). In addition, *Tfam* deletion resulted in a selective loss of mitochondrial gene expression at P10 (*mt-Nd1*, *mt-Co2*), while nuclear *Sdha* expression was unaffected (**Figure S6B**). We also observed a significant reduction in MTCO1 (complex IV subunit) protein levels in ileal sections of *Tfam*cKO at P14 (**Figures S6C and S6D**). These results validated our neonatal *Tfam*cKO model and were consistent with findings from other *Tfam*KO models^45,54,55^.

**Figure 6.**
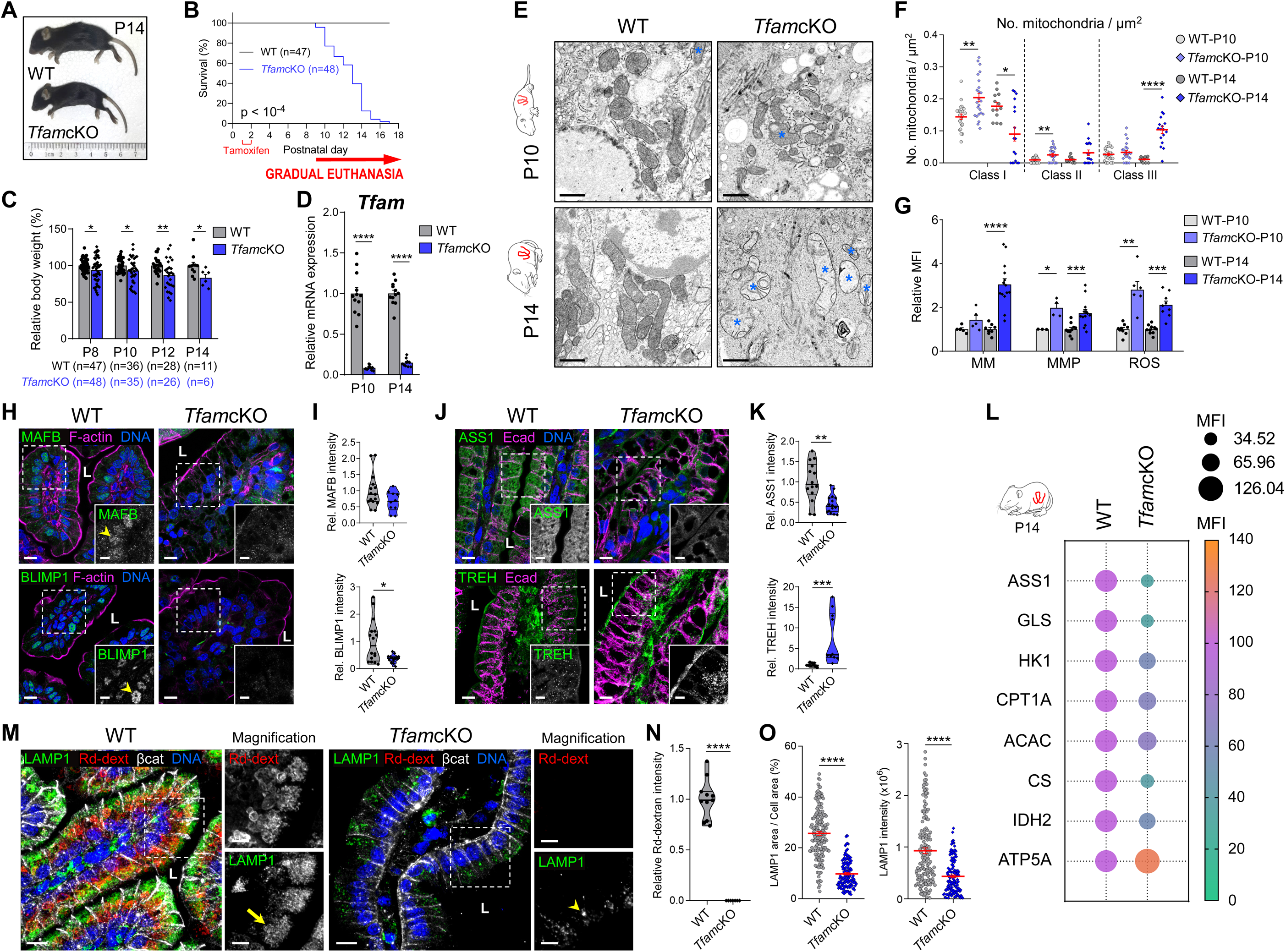
TfamcKO Neonates Recapitulate Endolysosomal and Metabolic Defects Observed in Atg16cKO Enterocytes, and Exhibit Weight Loss and Early Lethality. **(A)** Representative images of WT and *Tfam*cKO mice at postnatal day 14 (P14). **(B)** Kaplan–Meier survival curves of WT and *Tfam*cKO mice. Animals were euthanized if they failed to gain weight for two consecutive days or exhibited marked weight loss beginning at P9. **(C)** Relative body weight of WT and *Tfam*cKO mice at P8, P10, P12, and P14, expressed as percentage. Sample sizes are indicated at the bottom of the graph. **(D)** RT-qPCR analysis of *Tfam* mRNA in ileal epithelial cells from WT and *Tfam*cKO mice at P10 and P14 (n = 12,12,12,10). **(E)** *Tfam*cKO neonatal enterocytes show damaged mitochondria. TEM micrographs of ileal sections from WT (n = 3) and *Tfam*cKO (n = 4) mice at P10 and P14. Asterisks indicate damaged mitochondria. Scale bar, 1 μm. **(F)** Quantification of mitochondrial classes in (E), as defined in Figure 4B. Dots represent individual enterocytes. Sample sizes are indicated in Table S1. **(G)** Flow cytometry analysis of mitochondrial status of *Tfam*cKO neonatal enterocytes. Bar charts show relative MFI for Mitotracker Green (mitochondrial mass, MM), Mitotracker Red CMXROS (mitochondrial membrane potential, MMP) and CM-H_2_DCFDA (ROS) in ileal EpCAM^+^CD24^-^cKit^-^ cells from WT and *Tfam*cKO mice at P10 and P14. Sample sizes: MM (n = 5, 5, 8, 14), MMP (n = 3, 4, 10, 13), ROS (n = 9, 6, 10, 9). See Figure S3 for gating strategy. **(H)** Confocal images of ileal sections from WT and *Tfam*cKO mice at P14 (n = 3–4), stained with anti-MAFB (green, top) or anti-BLIMP1 (green, bottom), phalloidin (magenta), and DAPI (blue). Arrowheads mark nuclear localization of MAFB and BLIMP1 in WT LREs. **(I)** Violin plots quantifying nuclear MAFB (top; n = 15, 10) and BLIMP1 (bottom; n = 16, 16) intensity in (H), normalised to epithelial nuclear area. **(J)** Confocal images of ileal sections from WT and *Tfam*cKO mice at P14 (n = 3–4). Top: labelled with anti-ASS1 (green), anti-E-cadherin (Ecad, magenta), and DAPI (blue). Bottom: labelled with anti-TREH (green) and anti-Ecad (magenta). **(K)** Violin plots quantifying ASS1 (top; n = 14, 16) and TREH (bottom; n = 9, 10) intensity in (J), normalised to epithelial area. **(L)** Met-Flow profiling of ileal EpCAM^+^CD24^-^ cells from WT (n = 6) and *Tfam*cKO (n = 5) mice at P14. Bubble colour and size reflect group means for each marker. See Figure S3 for the gating strategy and Table S2 for statistical significance. **(M)** Confocal images of ileal sections from WT (n = 4) and *Tfam*cKO (n = 3) mice at P14 gavaged with rhodamine B–dextran (Rd-dext, red) 3 hours before sacrifice. Sections were stained with anti-LAMP1 (green), anti-β-catenin (white), and DAPI (blue). Arrow: endolysosomal system in WT LREs; arrowhead: reduced LAMP1⁺ vesicles in *Tfam*cKO enterocytes. Images deconvolved with Huygens software. **(N)** Violin plot quantifying Rd-dextran uptake (integrated density) in (M), normalised to epithelial area (n = 11, 7). **(O)** Quantification of lysosomal occupation and content in (M). Dot plots show the LAMP1⁺ area as a percentage of cell area (n = 167, 145) and LAMP1 intensity (integrated density) per enterocyte (n = 148, 139). In (D–O), mice aged P9–11 and P13–15 were grouped as “P10” and “P14,” respectively. In (H, J, M), dashed boxes indicate magnified areas; L, lumen. Scale bars: 10 μm (main); 5 μm (insets). In (C, D, G, I, K, L, N), data relative to WT values. In (C, D, G), graphs depict mean + SEM and dots represent individual mice. In (F, O), dot plots show mean ± SEM (red lines) and dots represent individual enterocytes. In (I, K, N), violin plots show median (thick line) and IQR (thin lines) and dots represent individual images (2–4 per mouse). Statistical analyses: Gehan–Breslow–Wilcoxon test (B), Student’s *t-*test (C, D, F [P14 Class III only], G, K, L, N), Mann–Whitney test (F [except P14 Class III], I, O). *p < 0.05, **p < 0.01, ***p < 0.001, ****p < 0.0001. **See also Figures S3, S6 and S7, and Tables S1 and S2.**

Further histological examination of the distal small intestine of *Tfam*cKO mice at P10 and P14 revealed substantial structural alterations. The epithelium exhibited clear signs of disorganization and inflammation, characterised by notably shorter, malformed villi, oedema, reduced GCs, and a thicker submucosa and lower mucosa (**Figures S6E-S6H**). Enterocytes also exhibited aberrant vacuoles, suggesting dysfunctions in nutrient processing (**Figure S6E**, arrows).

Moreover, flow cytometry analysis of the ileum at P10 and P14 revealed increased percentages of IIC (B cells, and Th cells) in *Tfam*cKO samples (**Figures S6I-S6K**). Given the elevated OXPHOS activity characteristic of intestinal stem cells (ISCs)^56^, we assessed ISC populations in ileal sections from P10 and P14 *Tfam*cKO mice via Olfm4 immunofluorescence (**Figure S6L**, arrows), which revealed a marked reduction in Olfm4⁺ crypts (**Figures S6M**). Additionally, lysozyme staining showed a striking increase in mislocalised PCs (**Figures S6N and S6O**). These PCs exhibited mislocalization, aberrant morphologies and enlarged granules, indicative of disrupted crypt architecture and compromised ISC niche homeostasis (**Figure S6N**, arrowheads).

With the systemic and intestinal phenotypes of *Tfam*cKO mice defined, we next investigated how loss of TFAM affects mitochondrial homeostasis in LREs. Firstly, mitochondria from *Tfam*cKO enterocytes at P10 and P14 were analyzed by TEM, following the same approach previously used in *Atg16*cKO mice. At P10, *Tfam*cKO enterocytes displayed more class I and II mitochondria, with reduced area and AR (**Figures 6E, 6F, and S6P**). By P14, damaged class III mitochondria increased while class I mitochondria decreased in number, size, shape and AR (**Figures 6E, 6F, and S6P**). This progressive mitochondrial damage resembles that observed in a previously described *Tfam*KO model in renal epithelial cells^55^. These data were corroborated by subsequent flow cytometry analysis, which revealed increased mitochondrial mass, membrane potential, and ROS levels in neonatal *Tfam*cKO enterocytes (**Figure 6G**), features reminiscent of those observed in TFAM-deficient lymphocytes^45,54^.

In summary, deletion of *Tfam* in the neonatal intestinal epithelium severely disrupted mucosal structure and function, leading to developmental defects that culminated in early mortality. Specifically, in LREs, TFAM loss caused the accumulation of damaged mitochondria and increased oxidative stress.

We next investigated whether mitochondrial dysfunction could similarly drive an accelerated metabolic transition in neonatal enterocytes, as observed in the *Atg16*cKO model. The microscopy analysis indicated a reduction in the expression levels of BLIMP1 and MAFB, with a more pronounced decrease in BLIMP1 (**Figures 6H and 6I**). As a result, we also observed lower levels of ASS1 and elevated expression of TREH in the *Tfam*cKO enterocytes (**Figures 6J and 6K**). These results were further supported by gene expression analyses of metabolic transition genes that are regulated by BLIMP1, MAFB, and c-MAF. We obtained a progressive downregulation of *Mafb*, *c-maf*, *Blimp1*, *Ass1,* and *Glb1*, accompanied by an upregulation of *Treh*, *Sis*, and *Arg2* at P14 (**Figure S7A**). Finally, the activity of key metabolic pathways was assessed in neonatal *Tfam*cKO enterocytes using Met-Flow^45,50^, revealing a marked reduction in nearly all metabolic proteins, except for ATP5A (**Figure 6L**), which was increased, possibly due to mitochondrial accumulation or as a compensatory response to mitochondrial dysfunction. Taken together, these findings indicate that neonatal *Tfam*cKO enterocytes undergo a premature metabolic transition, resembling the metabolic profile of PECs.

Following the analysis of mitochondrial defects in *Tfam*cKO neonatal enterocytes, we next investigated whether impaired mitochondrial function also perturbs the endolysosomal system of LREs. Although lysosome–mitochondria crosstalk has been predominantly characterized in the context of mitophagy, our findings suggest a reciprocal layer of regulation, whereby mitochondria control lysosomal biogenesis and function. This bidirectional communication implies that mitochondrial dysfunction not only disrupts energy metabolism but also secondarily compromises lysosomal integrity, thereby exacerbating the metabolic and endolysosomal defects observed in neonatal *Atg16*cKO enterocytes. Indeed, neonatal *Tfam*cKO enterocytes exhibited a complete loss of Rd-dextran uptake (**Figures 6M and 6N**) and a progressive downregulation of the endocytic genes *Dab2*, *Amn*, and *Mamdc4* (**Figures S7B-S7D**). Subsequently, we characterised a gradual reduction in the expression of lysosomal genes *Lamp2*, *Mcoln3*, and *Ctsl1* (**Figures S7E**), accompanied by a decrease in lysosomal content and size (**Figures 6M, 6O, S7F and S7G**). Thus, neonatal *Tfam*cKO enterocytes recapitulated the phenotype observed in the *Atg16*cKO model, validating a reciprocal interaction between mitochondria and lysosomes in LREs.

In conclusion, the *Tfam*cKO model enabled us to establish a bidirectional relationship between mitochondria and lysosomes by recapitulating the endolysosomal defects observed in autophagy-deficient neonatal enterocytes. Furthermore, we confirmed that mitochondrial and lysosomal dysfunction drive an early metabolic switch in LREs. These alterations, alongside epithelial inflammation, developmental delay, and premature death, position the *Tfam*cKO model as a valuable tool for investigating the pathophysiology of childhood malnutrition.

### Antioxidant activity in LREs modulates the onset of metabolic switch

After defining the metabolic switch underlying neonatal development, and the early metabolic switch in both *Atg16*cKO and *Tfam*cKO models, we focused on uncovering the mechanisms responsible for this phenotype. The observation that our gene-deficient mice presented heightened ROS levels in enterocytes (**Figures 4D and 6G**) suggested that redox dysregulation may underlie their accelerated metabolic transition. Indeed, the high mitophagic activity observed in LREs may contribute to the control of ROS^19,20^. Previous studies have shown that oxidative stress has been reported to repress *c-maf* expression^57^ and that antioxidant treatment can induce *Blimp1* expression^58^. Therefore, we hypothesise ROS regulation could represent a critical mechanism for controlling the timing of the metabolic transition in LREs.

We found that LREs exhibited significantly higher levels of mitochondrial ROS (mROS) compared to PECs (**Figure 7A**), possibly reflecting their elevated OXPHOS activity^59^. In contrast, total ROS levels were lower in LREs than in PECs (**Figure 7A**), indicating the presence of mechanisms that tightly regulate oxidative stress.

**Figure 7.**
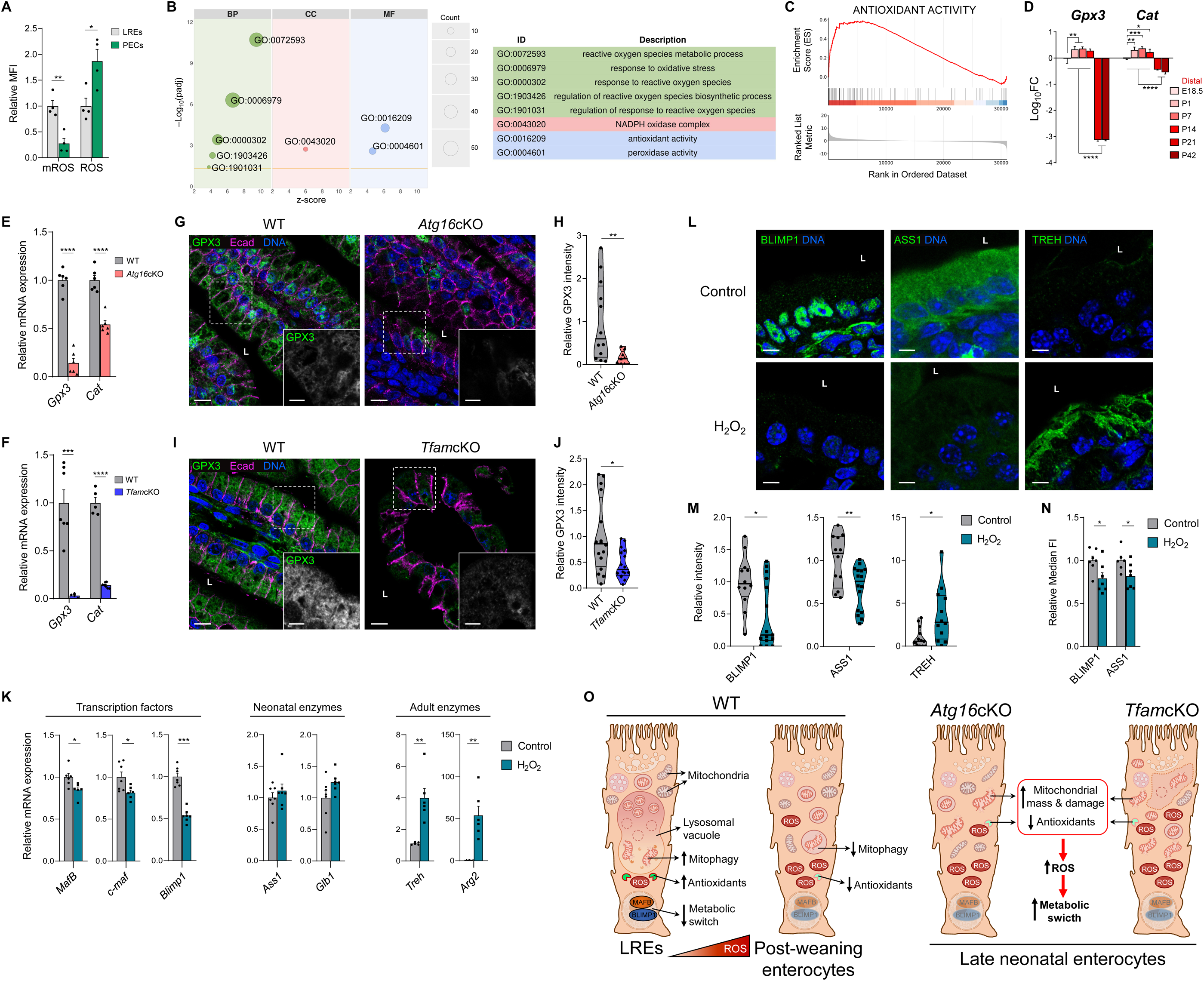
High Antioxidant Activity in LREs Regulates the Timing of Metabolic Transition. **(A)** Bar chart showing the relative mean fluorescence intensity (MFI) of MitoSOX (mitochondrial ROS) and CM-H_2_DCFDA (total ROS) in ileal EpCAM^+^CD24^-^ cells from WT mice at P7 (LREs) and P21–30 (PECs) (n = 4). Data relative to LREs values. See Figure S3 for gating strategy. **(B)** Bubble plot and list of ROS-related GO terms from over-representation analysis (ORA) of DEGs between LREs and PECs. The x-axis represents the z-score, the y-axis shows the significance as – log_10_(adjusted p-value), bubble size reflects the number of associated genes, and colours indicate GO categories (BP, biological process; CC, cellular component; MF, molecular function). **(C)** GSEA enrichment plot of antioxidant activity gene set (LREs vs PECs). **(D)** RT-qPCR analysis of *Gpx3* and *Cat* genes in ileal epithelial cells from WT mice at the indicated time points (n = 3). Values are expressed as log_10_ fold change relative to E18.5 embryonic levels. **(E)** RT-qPCR analysis of *Gpx3* and *Cat* genes in ileal epithelial cells from WT and *Atg16*cKO mice at P14 (n = 6). **(F)** RT-qPCR analysis of *Gpx3* (n = 7, 5) and *Cat* (n = 5, 7) in ileal epithelial cells from WT and *Tfam*cKO mice at P14. **(G)** Confocal images of ileal sections from WT and *Atg16*cKO mice at P14 (n = 3), stained with anti-GPX3 (green), anti-E-cadherin (Ecad, magenta), and DAPI (blue). **(H)** Violin plot of GPX3 intensity from (G), normalised to epithelial area (n = 12, 9). **(I)** Confocal images of ileal sections from WT and *Tfam*cKO mice at P14 (n = 4), labelled as in (F). **(J)** Violin plot of GPX3 intensity from (H), normalised to epithelial area (n = 16). **(K)** RT-qPCR analysis of transcription factor (*Mafb*, *c-maf*, *Blimp1*), neonatal (*Ass1*, *Glb1*) and adult (*Treh*, *Arg2*) enzyme genes in ileal epithelial cells from control and H2O2 treated mice at P12. Sample sizes: *Mafb* (n = 7, 6), *c-maf* (n = 7, 6), *Blimp1* (n = 7, 7), neonatal enzymes (n = 7, 7), *Treh* (n = 5, 6), *Arg2* (n = 6, 6). **(L)** Confocal images of ileal sections from control (n = 4) and H_2_O_2_-treated (n = 4–5) mice at P12, stained with anti-BLIMP1 (green, left) or anti-ASS1 (green, centre) or anti-TREH (green, right), and DAPI (blue). L, lumen. Scale bars, 10 μm. **(M)** Violin plots of BLIMP1, ASS1 and TREH intensity (integrated density) from (L). BLIMP1 data were normalised to epithelial nuclear area; ASS1 and TREH data, to epithelial area. Sample sizes: BLIMP1 (n = 12, 15), ASS1 (n = 12, 16), TREH (n = 13, 12). **(N)** Flow cytometry analysis of BLIMP1 (n = 7, 7) and ASS1 (n = 6, 7) in control and H_2_O_2_-treated mice at P12. Bar charts show relative median fluorescent intensity of BLIMP1 and ASS1 in ileal EpCAM^+^CD24^-^cKit^-^ cells. See Figure S3 for gating strategy. **(O)** Working model: In *Atg16*cKO and *Tfam*cKO neonatal enterocytes, mitochondrial dysfunction and reduced antioxidant activity elevate ROS levels, which repress *Blimp1* and *Mafb* expression and drive early metabolic transition. Enterocyte’s design inspired by Fujita et al.^9^. Figure created with BioRender.com. In (G, I), dashed boxes indicate magnified areas; L, lumen. Scale bars: 10 μm (main); 5 μm (insets). In (E, F, H, J), data relative to WT values; in (K, M, N), relative to control values. In (A, E, F, K, N), graphs represent mean + SEM and dots represent individual mice; in (D), mean + SD. In (H, J, M), violin plots show median (thick line) and IQR (thin lines) and dots represent individual images (2–4 per mouse, 3–5 mice per group). Statistical analyses: Student’s t test (A, E, F, H, K, M [ASS1], N), one-way ANOVA (D), Mann–Whitney test (J, M [BLIMP1, TREH]). *p < 0.05, **p < 0.01, ***p < 0.001, ****p < 0.0001. **See also Figure S3.**

Then, we further explored our LREs vs PECs RNA-Seq dataset analysis to identify additional pathways involved in ROS regulation. Numerous pathways related to ROS, ROS response, and antioxidant activity stood out (**Figure 7B and 7C**). Further, there was an overexpression of genes for the antioxidant enzymes Glutathione Peroxidase-3 (Gpx3) and Catalase (Cat) (not shown), suggesting that these antioxidants could be reducing ROS levels in LREs. Indeed, we observed that neonatal ileal samples showed induction of *Gpx3* and *Cat* expression compared to the embryo and post-weaning ileal samples, with the most drastic changes in *Gpx3* (**Figure 7D**). Moreover, in both *Atg16*cKO and *Tfam*cKO neonates, *Gpx3* and *Cat* expression were significantly decreased in the ileum (**Figure 7E and 7F**), and GPX3 protein levels were likewise reduced, as shown by IF (**Figure 7G-J**). Thus, oxidative stress may act as a trigger for the metabolic transition in LREs. To validate this hypothesis, neonatal mice were treated with hydrogen peroxide (H_2_O_2_) to induce oxidative stress^60^. mRNA expression analysis revealed a downregulation of the transcription factors *Blimp1*, *Mafb*, and *c-maf*, alongside an upregulation of adult enzymes such as *Treh* and *Arg2*. However, the expression of neonatal enzyme genes *Ass1* and *Glb1* remained unchanged (**Figure 7K**). Subsequently, we characterised a reduction of BLIMP1 and ASS1 levels by both IF and flow cytometry (**Figure 7L-7N**), while TREH expression was found to be increased (**Figure 7L and 7M**).

The data collectively suggest that elevated antioxidant activity is a key characteristic of LREs. When this antioxidant function is compromised in neonatal *Atg16*cKO and *Tfam*cKO enterocytes, alongside mitochondrial dysfunction, the oxyidative stress rise. This increase in ROS contributes to the repression of genes such as *Blimp1*, *Mafb*, and *c-maf*, ultimately triggering an early metabolic transition (**Figure 7O**). Therefore, this study emphasizses the critical role of mitochondrial-lysosomal coordination in mitigating oxidative stress and regulating the transcriptional programs essential for neonatal intestinal development. Additionally, it paves the way for future research into therapeutic strategies aimed at restoring mitochondria-lysosome homeostasis and redox balance in early gut disorders.

## DISCUSSION

In this work, we have elucidated new molecular mechanisms fundamental to the function and differentiation of mammalian LREs, highlighting the role of the mitochondria-lysosome axis. Disruption of autophagy in the neonatal gut, either through *Atg16*cKO or mitochondrial dysfunction via *Tfam*cKO, resulted in defective nutrient absorption and a reduction in lysosomal content. Most importantly, *Tfam*cKO neonates exhibited weight loss and premature death, resembling hallmarks of infant malnutrition. Notably, the early enzymatic maturation of *Atg16*cKO and *Tfam*cKO enterocytes indicates a direct connection between autophagic and mitochondrial processes in enterocytic maturation. Additionally, the associated increase in ROS observed in both models may contribute to this metabolic switch by suppressing the expression of transcription factors BLIMP1, MAFB, and c-MAF. These findings are consistent with our transcriptomic data, which reveal upregulation of mitophagy-and antioxidant-related pathways in LREs compared to PECs, supporting the concept that mitochondrial quality control and redox homeostasis are defining features of LRE identity and function. Taken together, our outcomes offer interesting mechanistic insights into the mitochondria-lysosome crosstalk and its role in postnatal intestinal physiology, with implications for understanding the pathophysiology of neonatal malnutrition and related disorders.

Autophagy and endocytosis are vesicle-mediated pathways that converge at the lysosome to degrade and recycle intra-and extracellular components. These processes intersect at multiple stages (vesicle formation, trafficking, and fusion) and share several key regulators. Mounting evidence indicates reciprocal dependence between these processes^25,61^. Using GFP-LC3 mice, we demonstrated that neonatal ileal samples exhibit increased autophagosome degradation compared to post-weaning counterparts (**Figures 1A-1C and S1C**). Consistently, *Atg16*cKO neonates displayed impaired absorption (**Figures 1H-1J**) and showed reduced expression of crucial endocytic genes characteristic of LREs (**Figures 2B-2F and S2G and S2H**). Moreover, *Atg16*cKO neonatal enterocytes displayed reduced lysosomal size and content (**Figures 2G-2J**), along with downregulation of key lysosomal genes (**Figures S2G and S2H**). These outcomes reveal a functional interplay between autophagy and the endolysosomal system in LREs.

Despite these endolysosomal defects, *Atg16*cKO mice reached the post-weaning stage without apparent physiological abnormalities, aside from mild signs of intestinal inflammation (**Figures S2A-S2D**). This observation aligns with findings in mice lacking the lysosomal cation channels Mucolipin-1 and -3 (Mcoln1 and Mcoln3), which also exhibit pronounced endolysosomal abnormalities in neonatal enterocytes but develop into phenotypically normal adults^12^. Together, these results suggest that endolysosomal and autophagic processes primarily affect LREs. Accordingly, once the neonatal period concludes and LREs are replaced by PECs, the phenotype becomes attenuated or fully resolves. However, further investigations will be needed to assess intestinal physiology and metabolism in adult *Atg16*cKO mice comprehensively.

We characterized a significant increase in autophagic activity in LREs as primarily due to the lysosomal degradation of mitochondria, a process known as mitophagy. Indeed, mitophagy-related genes exhibit a higher expression in neonatal ileum compared to post-weaning samples (**Figures 3B-3F**). Correspondingly, neonatal *mito*-QC mice^42^ demonstrated enhanced mitochondrial degradation within the LVs of LREs compared to PECs (**Figures 3H and 3I**). Furthermore, this process was inhibited either by leupeptin treatment^43^ (**Figures 3H and 3I**) or through genetic ablation of autophagy (*Atg16*cKO) (**Figures 3L and 3M**), confirming the dependence of mitochondrial clearance on lysosomal-autophagic activity. Consistent with this, neonatal *Atg16*cKO enterocytes accumulated dysfunctional mitochondrial mass (**Figure 4**).

Specifically, these cells showed an increase in the area of damaged mitochondria (class III), whereas healthy mitochondria (class I) exhibited a smaller area but an increased aspect ratio (**Figures 4A-4F**). These differences possibly reflect not only mitophagy blockade but also alterations in mitochondrial dynamics, such as enhanced fusion or reduced fission, which are aimed at preserving oxidative phosphorylation (OXPHOS) capacity^62^. Future studies should investigate the expression of proteins involved in mitochondrial fusion and fission in the *Atg16*cKO model to define better the molecular landscape that integrates mitochondrial dynamics with mitophagy in LREs.

After identifying mitochondrial homeostasis defects secondary to impaired endolysosomal function in the *Atg16*cKO model, we next investigated the reverse relationship by directly disrupting mitochondrial function using the *Tfam*cKO model. *Tfam*cKO neonates exhibited severe growth impairment and premature death (**Figures 6A-6C**), and their enterocytes accumulated markedly damaged mitochondria (**Figures 6E-6G and S6P**). These mice also reproduced key features of the *Atg16*cKO phenotype, including loss of uptake capacity, reduced lysosomal content, and downregulation of endolysosomal genes (**Figures 6M-6O and S7B-S7G**). Interestingly, these lysosomal defects differ from what has been observed in *Tfam*-deficient T and B lymphocytes, where dysfunctional acidic lysosomal compartments are expanded^45,54^. In neonatal enterocytes, however, Tfam deletion results in diminished lysosomal size and content, highlighting that mitochondria–lysosome interactions are highly cell-type specific. Beyond this bidirectional crosstalk, our data raise the possibility of a retrograde signalling mechanism. Mitochondria are increasingly recognized as signalling hubs that influence nuclear transcriptional programs, as shown in immune cells. In LREs, two interconnected processes may be at play: first, endolysosomal-mediated nutrient internalization, which is defective in both *Atg16*cKO and *Tfam*cKO models; and second, mitophagy-driven mitochondrial degradation, which provides retrograde signals to the nucleus. This latter pathway could be essential for controlling the transcriptional networks that govern the metabolic switch (via BLIMP1 and MAFB) and for sustaining lysosomal functionality itself. In this view, lysosomes must degrade mitochondria not only to preserve organelle quality control but also to control the transcriptional reprogramming required for the neonatal-to-adult metabolic transition.

At weaning, the mammalian intestine undergoes a shift to extracellular nutrient digestion and adjusts its enzymatic profile to meet the metabolic demands of a carbohydrate-rich solid diet^3^. At the same time, LREs are replaced by PECs lacking large lysosomal vacuoles^46^. The transcription factors BLIMP1 (PRDM1), along with MAFB and c-MAF (both members of the Maf protein family) have been positioned as central repressors of this switch. Neonatal mice lacking these transcription factors in the intestinal epithelium exhibited impaired nutrient absorption, early downregulation of neonatal enzymes, and upregulation of adult counterparts, ultimately leading to developmental defects^47–49^. Furthermore, our research demonstrated that dysfunctions in autophagic, lysosomal, and mitochondrial processes in LREs —induced by the deletion of *Atg16l1* and *Tfam*— led to a repression of these transcription factors, resulting in similar phenotypes (**Figures 1H, 5, 6H-6N and S7A)**.

We further analysed the activity of core metabolic pathways in WT, *Atg16*cKO, and *Tfam*cKO neonatal enterocytes, comparing them to PECs (**Figures 5H, 6L and S5E)**. This analysis revealed a moderate reduction in metabolic activity in *Atg16*cKO cells and a more noticeable decline in *Tfam*cKO enterocytes, suggesting a shift toward an adult-like metabolic state. These findings contrast with previous transcriptomic data from neonatal mice lacking MAFB and c-MAF, which reported an upregulation of OXPHOS and fatty acid degradation pathways^47^. However, that study did not include a direct comparison to PECs, leaving open the possibility of compensatory mechanisms. In this context, our results may provide a more accurate reflection of the metabolic trajectory during neonatal-to-adult transition and its dependence on the mitochondria-lysosome axis.

We have defined the metabolic evolution of enterocytes during neonatal development beyond the genes classically associated with the transcription factors BLIMP1, MAFB, and c-MAF, and established a functional link between this transition and the maintenance of lysosomal and mitochondrial homeostasis. At the same time, our data reinforce the idea that, in enterocytes, genetic programs prevail over the sensing of external cues: despite the neonatal context (milk-based diet, reduced microbiota, and limited immune activation)^63^, the repression of these transcription factors leads to inevitable epithelial maturation. Yet, the cellular mechanisms that regulate these genetic pathways still need to be clarified.

Reactive oxygen species (ROS) are oxygen-derived molecules that play a crucial role in various cellular processes, including energy production, mitochondrial signalling, and cell differentiation^64,65^. Their intracellular levels are tightly regulated through multiple mechanisms: the mitochondrial electron transport chain constitutes a major source of ROS, while antioxidant enzymes serve as key systems for neutralizing their excess^59,64,65^. In the intestine, ROS have been primarily associated with more differentiated epithelial cells exhibiting higher OXPHOS activity^24^. However, their role in epithelial maturation during neonatal intestinal development continues to be unexplored.

Our results indicate that LREs display a transcriptional signature consistent with elevated antioxidant activity, possibly as a compensatory mechanism to counteract high levels of mitochondrial ROS resulting from hyperactivated OXPHOS (**Figures 7A-7D**). Beyond this homeostatic role, recent studies suggest that this antioxidant signature may also be linked to the repression of transcription factor genes *Blimp1*, *Mafb*, and *c-maf*, and consequently, to the maintenance of LRE identity and function. The antioxidant vitamin ascorbic acid has been shown to increase the expression of Blimp1 in B lymphocytes through the activity of DNA demethylases TET2 and TET3^58^. Conversely, treatment with H_2_O_2_ decreases the expression of *c-maf*, whereas the ROS scavenger N-acetyl cysteine (NAC) can restore its levels in human adipose tissue-derived mesenchymal stem cells^57^.

This hypothesis is further supported by the elevated ROS levels and the loss of the antioxidant signature observed in our knockout models (**Figures 4G, 6G, 7E-7J)**, as well as by our experiments in which H_2_O_2_ treatment of neonatal WT mice accelerated the metabolic transition of enterocytes (**Figures 7K-7N**). Together, these outcomes suggest that both ROS levels and ROS-regulated genetic programs actively shape epithelial differentiation and metabolic programming in the neonatal intestine (**Figures 7O**). This may represent a physiological mechanism to prevent premature transitions until the tissue is developmentally prepared. In this context, therapeutic strategies aimed at fine-tuning redox homeostasis could offer a powerful approach to supporting neonatal intestinal health.

In addition to their role in nutrient absorption, LREs may act as immunometabolic sentinels during the neonatal period, integrating microbial cues and intracellular metabolic status to modulate immune responses. Due to their high endolysosomal activity, LREs could internalise and process luminal antigens, including microbial components, thereby contributing to the development of the intestinal immune system^66^. In line with this, our RNA-Seq analysis revealed multiple differentially regulated immune-related pathways (data not shown).

Disruption of lysosomal and mitochondrial homeostasis in *Atg16*cKO and *Tfam*cKO mice likely impairs these immune functions, as suggested by the inflammatory status observed in both models (**Figures S2D-S2F, S6E, and S6H-S6K**). In fact, mice with autophagy defects also exhibit gut microbiota alterations and immune dysregulation^17^. Furthermore, cytosolic mtDNA resulting from mitochondrial damage due to TFAM deficiency has been shown to activate the cGAS–STING inflammatory pathway^55^, while mitophagy induction, along with improved mitochondrial status, attenuates this response^43^. Additionally, *c-maf* KO mice exhibit changes in intraepithelial lymphocytes and microbiota composition^67^, suggesting that genetic programs involved in epithelial maturation, affected in our KO models, may be tightly linked to intestinal immune and microbial homeostasis.

Given the immunometabolic imbalance observed in *Atg16*cKO and *Tfam*cKO neonates, it is plausible that these models experience alterations in their gut microbiota. These changes may not only reflect enterocytic dysfunction but could also actively contribute to the phenotype by disrupting microbial-derived cues essential for LRE maintenance and programming. Future studies involving microbiota profiling and transplantation will help determine whether restoring a balanced microbial ecosystem can rescue LRE function.

Finally, the potential microbiota alterations described above, together with the inflammation and oxidative stress observed in our KO models, suggest parallels with known hallmarks of children’s malnutrition disorders. Particularly, *Tfam*cKO mice display a more pronounced phenotype and marked developmental defects, resembling features of Kwashiorkor, a severe form of protein deficiency prevalent in low-income settings ^68,69^. This redox imbalance—marked by low antioxidant levels and increased ROS—in our KO models closely mirrors that seen in malnourished children, and may contribute to the broader clinical phenotype of undernutrition^68^. Accordingly, pilot studies have demonstrated that antioxidant supplementation may provide therapeutic benefit in conditions such as Kwashiorkor^70^. Moreover, our H_2_O_2_ treatment experiments, which replicated the epithelial maturation phenotype characterized in our gene-deficient mice, might similarly induce nutritional, microbial, and inflammatory disturbances. Indeed, intestinal tissues from H_2_O_2_-treated neonates displayed signs of inflammation (data not shown).

Collectively, these considerations suggest that malnourished children may also experience premature intestinal maturation and highlight the need for deeper investigation into antioxidant therapies as potential interventions. Furthermore, our KO models may serve as valuable experimental platforms to investigate the mechanisms underlying neonatal malnutrition and to explore interesting avenues for its treatment. Altogether, future studies aimed at restoring LRE mitophagy and metabolism, as well as modulating the redox balance, could yield promising strategies to counteract the devastating consequences of malnutrition-related disorders in early life.

In conclusion, our study uncovers novel mechanisms underlying the function and metabolic transition of neonatal enterocytes. We demonstrate that lysosomal and mitochondrial homeostasis is indispensable for efficient nutrient absorption and for sustaining the high metabolic activity required during this energy-demanding stage of life. Further investigation of the mitochondria-lysosome axis in LREs will not only advance our understanding of neonatal intestinal physiology but also contribute significantly to the development of targeted interventions against childhood malnutrition.

### Limitations of the study

This work demonstrates that neonatal intestinal deficits in autophagy and mitochondrial homeostasis lead to endolysosomal dysfunction in enterocytes, impairing development and ultimately triggering inflammation and early lethality. However, since we used *Villin-cre*-based models, where gene deletion occurs in all intestinal epithelial cells, we cannot exclude their contribution to the observed phenotypes. Indeed, we detected altered abundance and morphology of these cells in both *Atg16*cKO (**Figures 1F, 1G, and S2E-S2F**) and *Tfam*cKO mice (**Figures S6G, S6L-S6O**). Previous studies support this interpretation: defects in GCs and PCs are linked to intestinal inflammatory diseases^71,72^, and intestinal autophagy loss has been associated with increased oxidative stress in ISCs^36^, which could partially explain the elevated ROS levels in *Atg16*cKO enterocytes (**Figure 4D**). Similarly, *Tfam* deletion in the adult intestinal epithelium causes ISC loss and lethality^53^, suggesting that depletion of ISCs may contribute to the early death of neonatal *Tfam*cKO mice, limiting our ability to study mitochondrial function in enterocytes at later developmental stages.

To address these limitations, we plan to develop an LRE-specific tamoxifen-induced conditional knockout model to target critical genes specifically in neonatal enterocytes. Although selectively targeting neonatal enterocytes would yield a more precise model, this is hindered by the lack of fully specific promoters and the technical difficulty of isolating and culturing LREs, which are highly sensitive to dissociation-induced cell death. To overcome these challenges, we are also studying *in vitro* systems that recapitulate the properties of LREs and may serve as a platform for future functional validation.

## RESOURCE AVAILABILITY

### Lead contact

Further information and requests for resources and reagents should be directed to and will be fulfilled by the lead contact, Fernando Martin-Belmonte.

### Materials availability

This study did not generate new unique reagents.

### Data and code availability

RNA-seq Illumina paired-end reads (FASTQ) generated for this study are available at the European Nucleotide Archive (ENA; http://www.ebi.ac.uk/ena/) under the study accession number PRJEB46511.

## ACKNOWLEDGMENTS

We thank Carmen M. Ruiz-Jarabo for her comments on the manuscript and members of the Martin-Belmonte and Martinez-Martin labs for their support and helpful discussions. This work was supported by MICINN (BFU2017-83243-P; PID2020-120367GB-I00; PID2023-151844OB-I00; RED2022-134927-T), Fondos FEDER investigación (Horizon-MSCA-2022-DN-101119504) and Fundación Ramon Areces (CIVP18A3904). HR22-00447 from “La Caix”, RTI2018-101586-A-I00, PID2021-126298OB-100 to N.M.-M. G.H. and M.I.-P. were supported by FPI fellowships BES-2018-077789 and PRE2019-089076, respectively, funded by the Spanish Ministry of Science and Innovation and the State Research Agency (MCIN/AEI /10.13039/501100011033). D.A.L was supported by FPU and JAE fellowships FPU23/03569 and JAEINT_22_01549. L.A was supported by Horizon-MSCA-2022-DN -101119504. I.G.G. was funded by the Medical Research Council, UK (MC_UU_00038/2). We thank Victor Lazaro Morato (currently at IMN-CNM, CSIC, Madrid, Spain) for his contribution to the initial analyses of the general phenotype and mitochondrial status by TEM in the *Tfam*cKO model. We acknowledge Michel Bagnat (Duke University, USA) for the generous donation of mCherry plasmid, and Juan M. Serrador, Catalina Grabowski and Dionisio Ureña (CBM), for their essential contribution to the protocol for the growth and purification of mCherry plasmid. We are thankful to the Animal, Flow Cytometry, Advanced Optical Microscopy, and Electron microscopy facilities at the CBM, as well as the Histology Service at the Centro Nacional de Biotecnologia (CNB, CSIC, Madrid, Spain), for their excellent technical support. We acknowledge the Centro de Análisis Genómico (CNAG-CRG, Barcelona, Spain) for the RNA sequencing, and to Biocomputational Analysis Core Facility at the CBM, specially to Sandra González de la Fuente, for the NGS data analysis and plotting. We thank the computational resources and support provided by the Centro de Computación Científica at Universidad Autónoma de Madrid (CCC-UAM).

Certain Figures were inspired by or adapted from previously published sources, some of which are licensed under Creative Commons (CC BY 4.0) (https://creativecommons.org/licenses/by/4.0/). While every effort has been made to credit original sources, any omissions are unintentional and will be corrected in future revisions if identified. Some figures were created with BioRender.com.

## SUPPLEMENTAL INFORMATION

**Figure S1.**
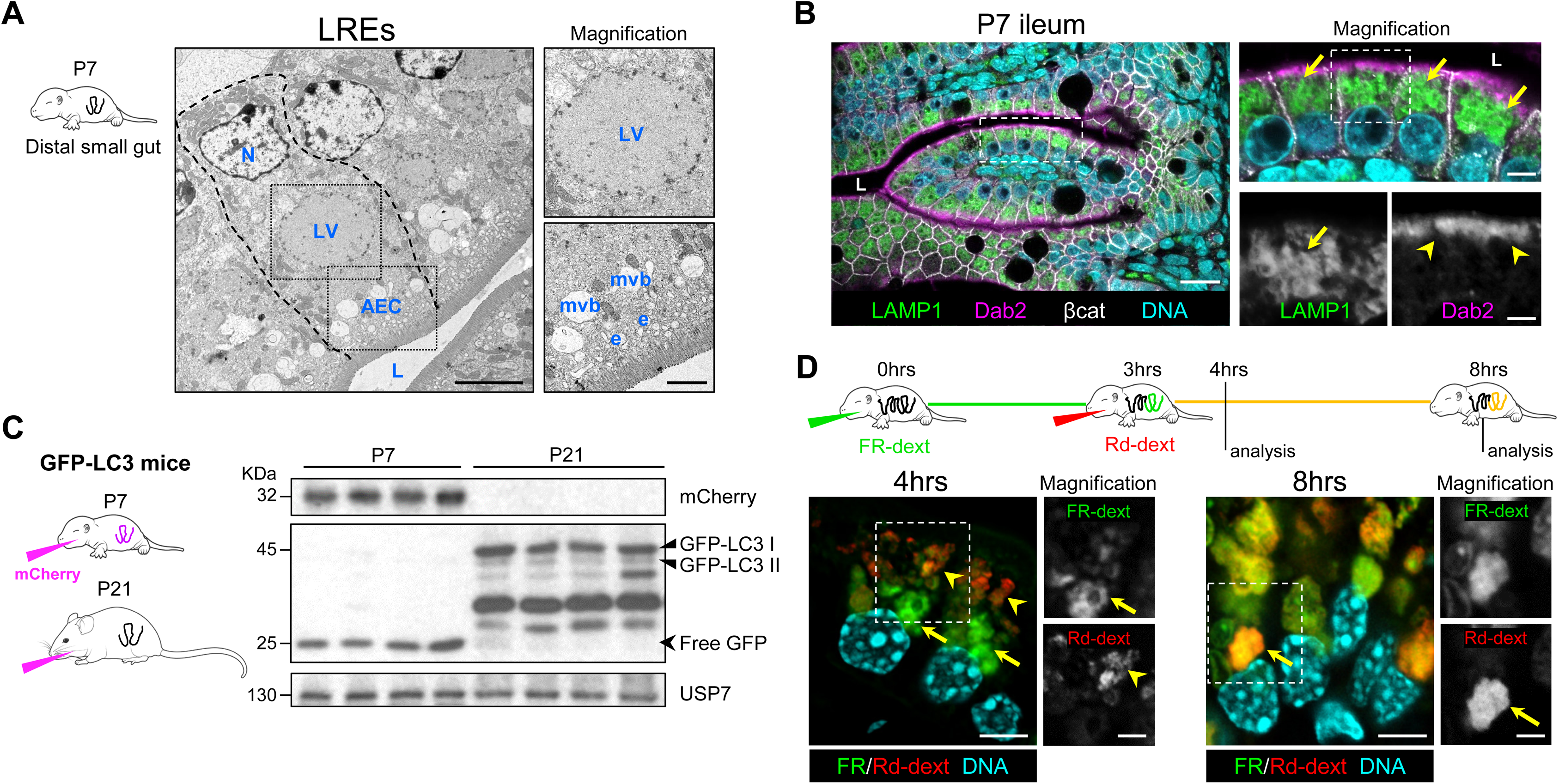
Characterization of LREs in the Neonatal Mouse Ileum. **Related to Figures 1 and 2** (A) Representative TEM image of LREs from the distal small intestine of a P7 mice (n = 3). The profile of an LRE is outlined by a dashed line. Dashed boxes indicate magnified areas. N, nucleus; LV, lysosomal vacuole; AEC, apical endosomal complex; e, endosome; mvb, multivesicular body; L, lumen. Scale bars: 5 μm (main); 2 μm (insets). (B) Confocal microscopy images of ileal sections from P7 mice (n = 3), stained with anti-LAMP1 (green), anti-Dab2 (magenta), anti-β-catenin (white), and DAPI (cyan). Dashed boxes highlight magnified areas. Arrows indicate the endolysosomal system; arrowheads point to apical Dab2 localization. Scale bars: 20 μm (main); 10 μm (first magnification); 5 μm (second magnification). (C) Western blot of mCherry and GFP-LC3 isoforms in ileal epithelial cells from GFP-LC3 mice at P7 and P21, gavaged with mCherry 3 hours before sacrifice. USP7, loading control. (D) Sequential fluorescent dextran uptake assay. P7 mice were gavaged with Alexa 647-dextran (FR-dext, green), followed by rhodamine B–dextran (Rd-dext, red) 3 hours later. Mice were sacrificed at 4 and 8 hours post-initial gavage to assess uptake via confocal microscopy (n = 3). Arrows point to lysosomal vacuoles containing both dextrans at 8 hours. Arrowheads indicate endosomes. Scale bars: 5 μm (main); 2 μm (insets).

**Figure S2.**
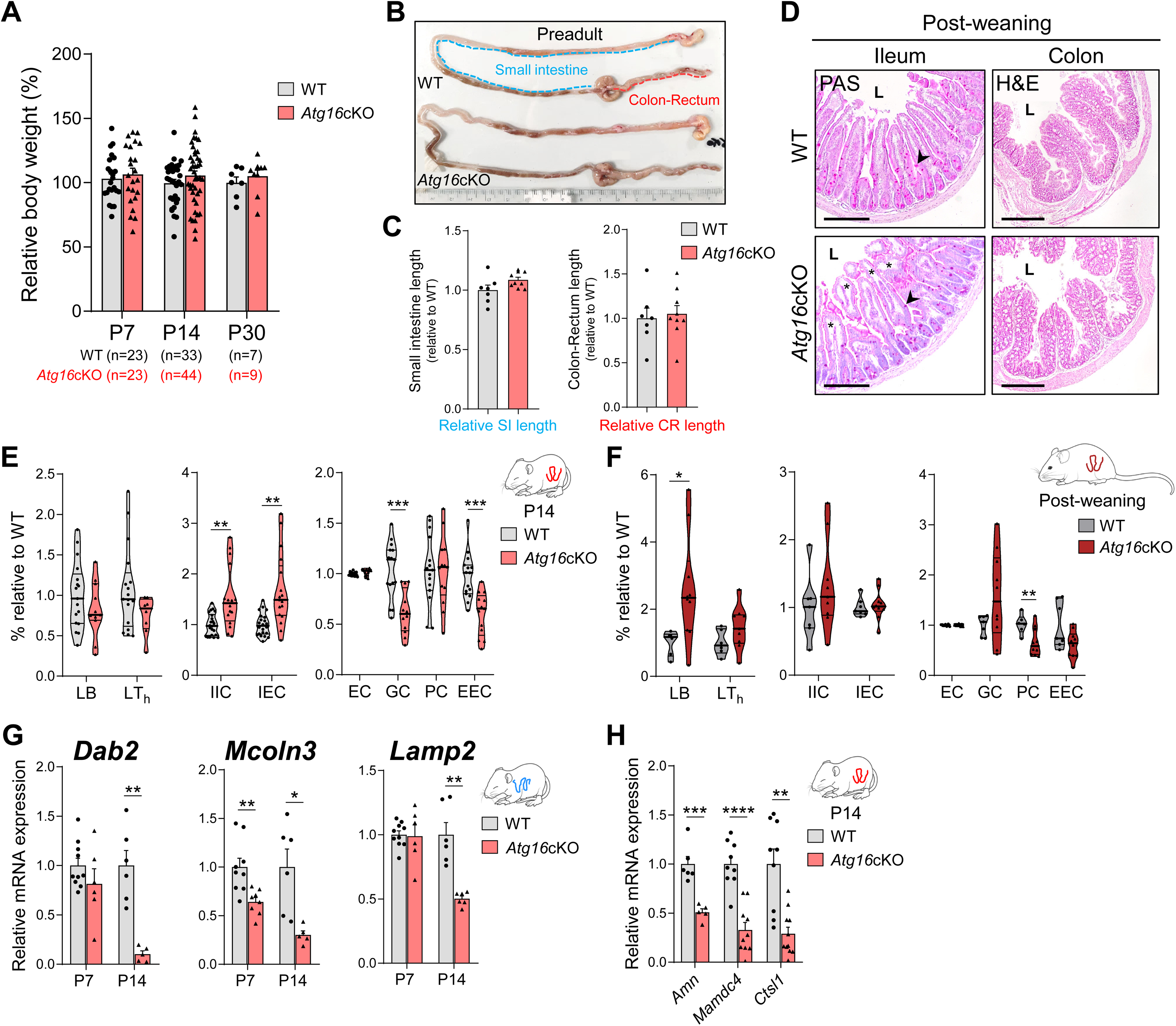
Phenotype of Atg16cKO Mice. **Related to Figures 1 and 2** **(A)** Relative body weight of WT and *Atg16*cKO mice at postnatal days P7, P14, and P30. Sample sizes are indicated in the graph. **(B)** Representative images of intestines from WT and *Atg16*cKO mice at the post-weaning stage (P30). **(C)** Quantification of intestinal lengths shown in (B). Bar graphs show the relative lengths of the small intestine (SI) and the colon–rectum (CR) segments (n = 7, 9). **(D)** Representative histological images of ileum and colon sections from WT and *Atg16*cKO post-weaning (P30) mice (n = 3). Ileal sections are stained with periodic acid-Schiff (PAS) and haematoxylin; colon sections are stained with haematoxylin and eosin (H&E). Asterisks denote oedema in *Atg16*cKO ileum; arrowheads indicate goblet cells. Scale bars: 200 μm (ileum), 400 μm (colon). **(E and F)** Flow cytometry analysis of immune and epithelial cell populations in the ileum of WT and *Atg16*cKO mice at P14 (E) and P30 (F). Left: B lymphocytes (LB) and T-helper lymphocytes (LTh) percentages. Centre: Innate immune cells (IIC) and intestinal epithelial cells (IEC) percentages. Right: Enterocyte and enterocytic progenitors (EC), goblet cell (GC), Paneth cell (PC), and enteroendocrine cell (EEC) percentages. See Figure S3 for gating strategy. (E): Left, n = 17,8–11; Centre, n = 22,16–17; Right, n = 15,12–13. (F): n = 6–7, 9–10. **(G)** RT-qPCR analysis of *Dab2* (n = 10,6,6,5), *Mcoln3* (n = 9,8,6,5), and *Lamp2* (n = 10,6,6,6) in epithelial cells from the proximal small intestine of WT and *Atg16*cKO mice at P7 and P14. **(H)** RT-qPCR analysis of *Amn* (n = 6,5), *Mamdc4* (n = 9,10), and *Ctsl1* (n = 9,11) in ileal epithelial cells from WT and *Atg16*cKO mice at P14. In (A, C, G, H), graphs represent mean + SEM. In (E, F), violin plots depict median (thick line) and interquartile range (IQR, thin lines). In all graphs, data relative to WT values, dots represent individual mice and Student’s t test was conducted. *p < 0.05, **p < 0.01, ***p < 0.001, ****p < 0.0001.

**Figure S3.**
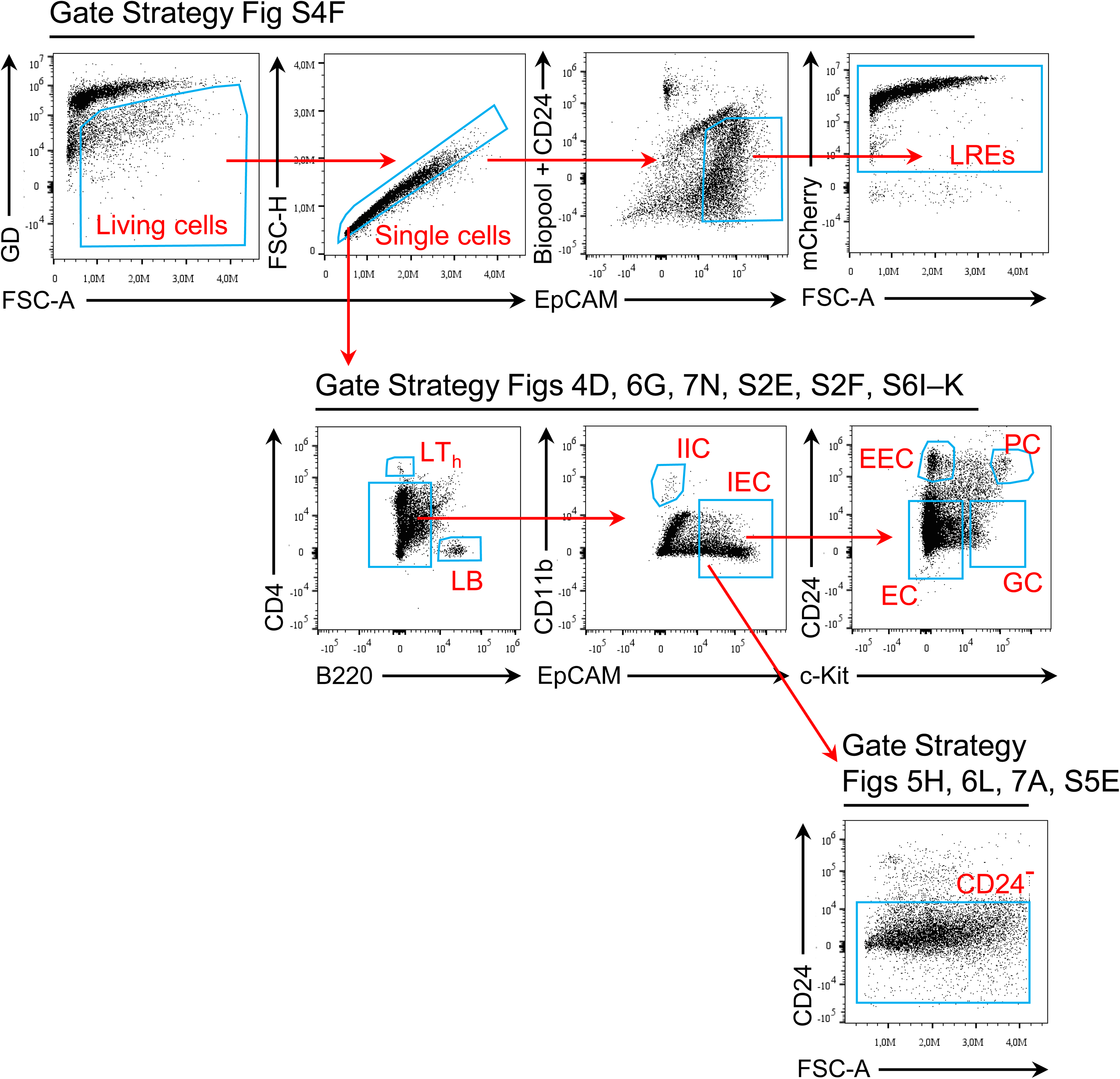
Gating Strategies Used for Cell Population Characterization by Flow Cytometry. **Related to Figures 4–7, S2 and S4–S6** Biopool is the combination of immune cell markers anti-CD11c, anti-CD3e, anti-CD11b, anti-CD19, and anti-CD45.2. The strategy for selecting IEC subtypes is based on Asano et al. ^36^. GD, ghost dye; LT_h_, T-helper lymphocytes; LB, B lymphocytes; IIC, innate immune cells; IEC, intestinal epithelial cells; EEC, enteroendocrine cells; PC, Paneth cells; EC, enterocytes and enterocytic progenitors; GC, goblet cells.

**Figure S4.**
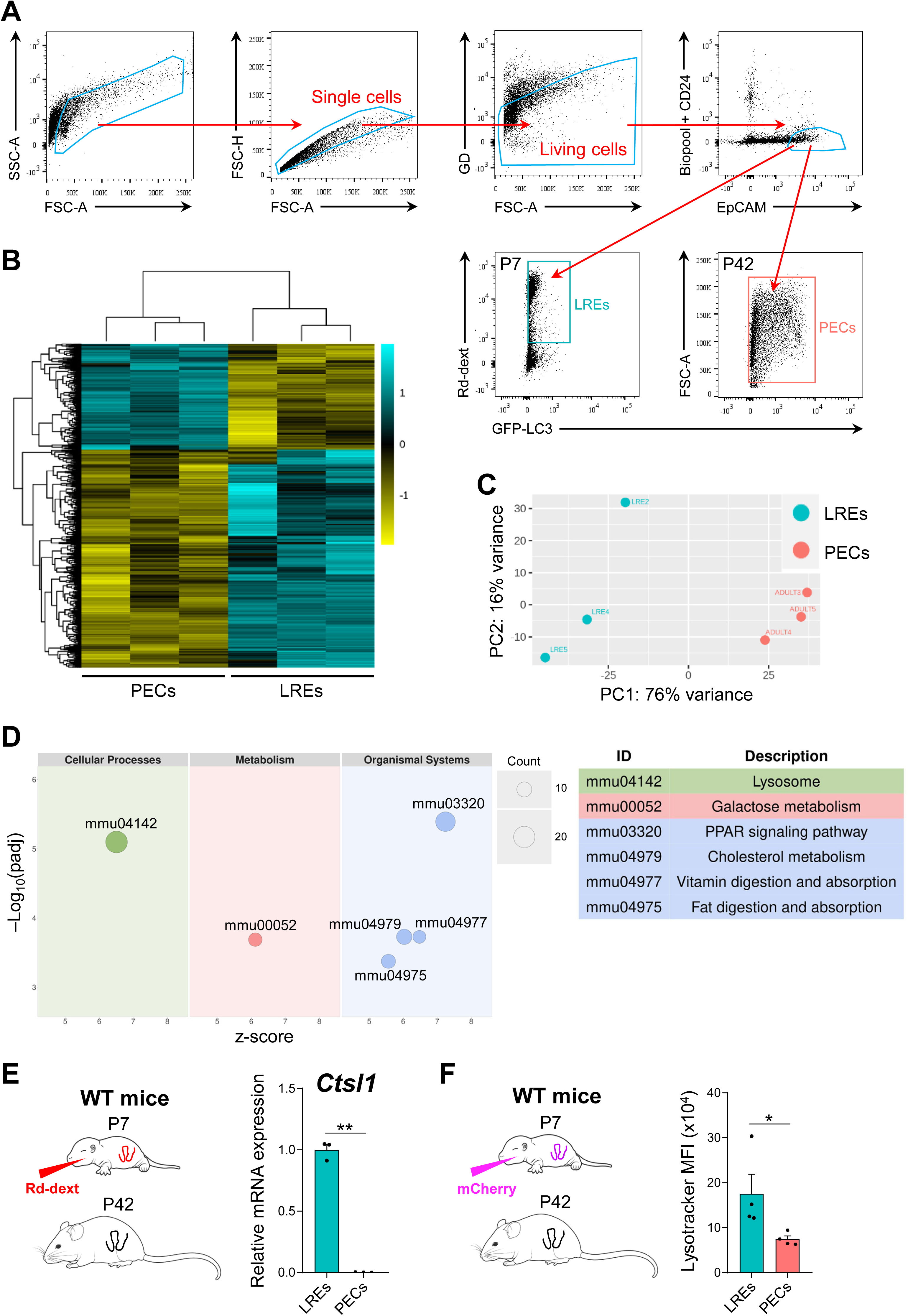
RNA-seq Analysis Comparing Gene Expression of LREs and PECs. **Related to Figure 3** (A) FACS gating strategy used to isolate LREs and PECs from GFP-LC3 mice at P7 (gavaged with Rd-dextran, 16 hours) and P42, respectively, for RNA-seq analysis. Single-living EpCAM^+^CD24^-^Biopool^-^ cells were gated. For LREs, Rd-dext⁺GFP^high^ cells were selected. (B) Heatmap with hierarchical clustering of differentially expressed genes (DEGs) with q-value < 0.05 reveals distinct expression patterns between LREs and PECs. Each row represents a single gene and each column represents a condition sample. Each sub-column represents an individual sample (3 samples per group). Gene expression intensities were log_2_-transformed and visualized as a colour gradient from yellow to light blue, representing the shift from downregulation to upregulation, as indicated in the key. (C) Principal component analysis (PCA) plotting for 2 groups based on normalised overall transcriptomic reads. The PCA figure represents a two-dimensional scatterplot of the first two principal components of the RNA-Seq data. Analyses in panels (B) and (C) show that replicates of the same cell population are more similar to each other than to replicates from a different population. (D) Bubble plot of selected Kyoto Encyclopaedia of Genes and Genomes (KEGG) pathways from over-representation analysis (ORA) of the differential gene expression data from the LREs vs PECs. The x-axis represents the z-score, the y-axis shows the significance as –log10(adjusted p-value), bubble size reflects the number of associated genes, and colours indicate KEGG pathway categories. (E) RT-qPCR validation of *Ctsl1* expression in LREs and PECs from WT mice at P7 (gavaged with Rd-dextran, 16 hours) and P42, respectively. FACS was used for cell sorting, selecting single-living EpCAM^+^CD24^-^Biopool^-^ cells, with Rd-dext⁺ cells further gated for LREs. In LREs, each dot represents one of three pooled samples (n = 3), each composed of multiple mice; in PECs, each dot corresponds to an individual mouse (n = 3). Data relative to LREs values. (F) Analysis of lysosomal content in LREs and PECs by measuring Lysotracker MFI by flow cytometry. Bar chart depicts Lysotracker MFI of LREs and PECs from WT mice at P7 (gavaged with mCherry, 16 hours) and P42 respectively. Single-living EpCAM^+^CD24^-^Biopool^-^ cells were gated. For LREs, mCherry⁺ cells were selected. See Figure S3 for gating strategy. Dots represent individual mice (n = 3). Graphs in (E, F) represent mean + SEM. In (E), Student’s t test; in (F), Mann–Whitney test. *p < 0.05, **p < 0.01.

**Figure S5.**
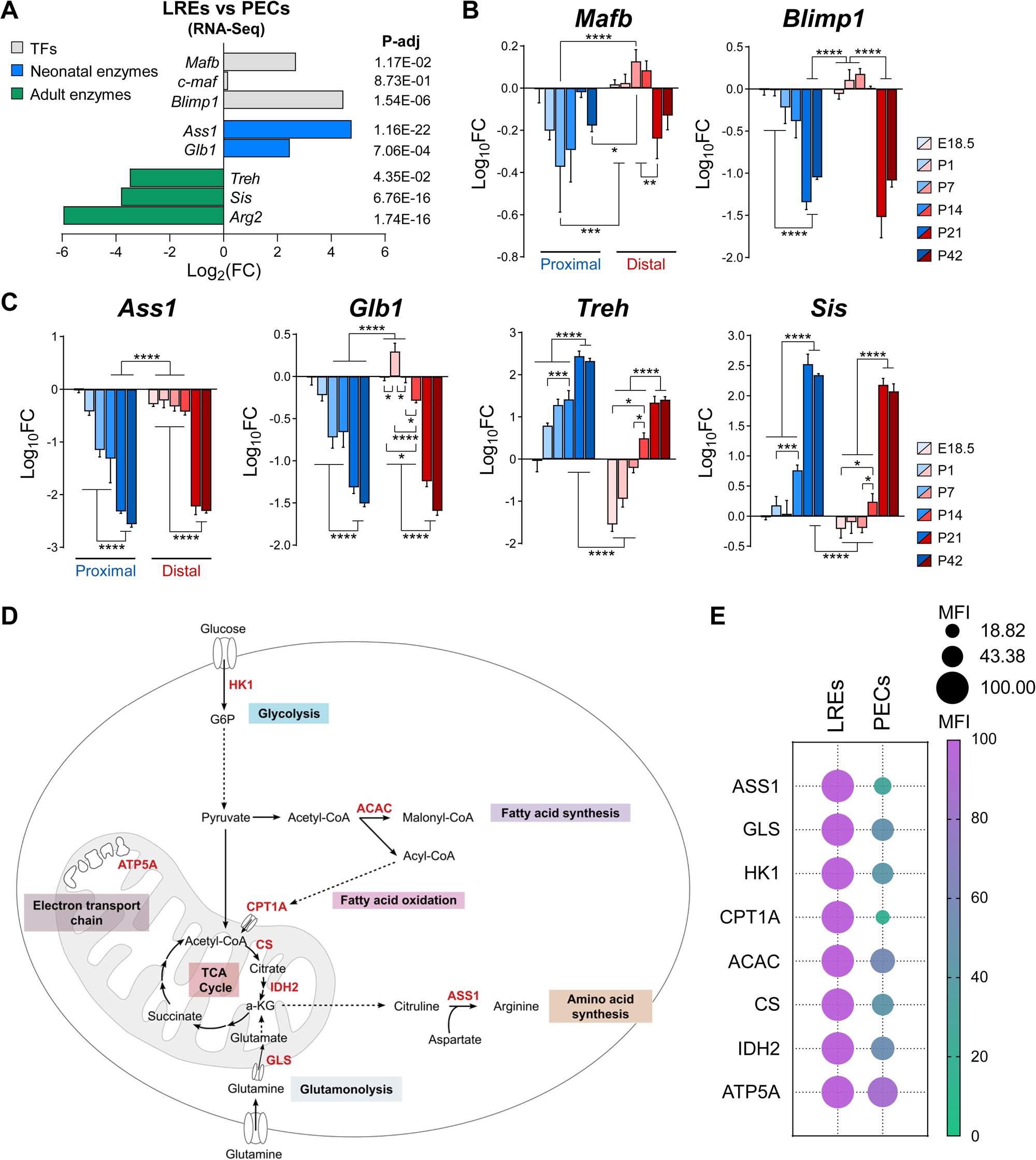
Metabolic Transition in the Small Gut During Postnatal Development. **Related to Figure 5** (A) Expression and statistical data of selected genes encoding transcription factors (TFs: *Mafb, c-maf, Blimp1*), neonatal enzymes (*Ass1, Glb1*), and adult enzymes (*Treh, Sis, Arg2*) from the LREs vs PECs RNA-seq analysis. **(B and C)** RT-qPCR analysis of *Mafb* and *Blimp1* (B), and of *Ass1, Glb1, Treh,* and *Sis* (C) genes in epithelial cells isolated from proximal and distal small intestinal segments of WT mice at the indicated postnatal time points (n = 3). Data are normalised to the expression levels in the proximal small intestine of E18.5 embryos and expressed as log_10_fold change (log_10_FC). **(D)** Schematic representation of the cellular metabolic pathways. Enzymes used in the Met-flow panel are shown in red. Scheme adapted from Iborra et al. ^45^ (CC BY 4.0). **(E)** Met-Flow analysis of ileal EpCAM^+^CD24^-^ cells from WT mice at P7 (LREs) and P42 (PECs) (n = 2–4). Data relative to LREs values. Bubble colour and size reflect group means for each marker. See Figure S3 for the gating strategy and Table S2 for statistical significance. In (B, C), graphs represent mean + SD, two-way ANOVA was conducted and only the most relevant statistically significant comparisons are shown. In (E), Student’s *t-*test. *p < 0.05, **p < 0.01, ***p < 0.001, ****p < 0.0001.

**Figure S6.**
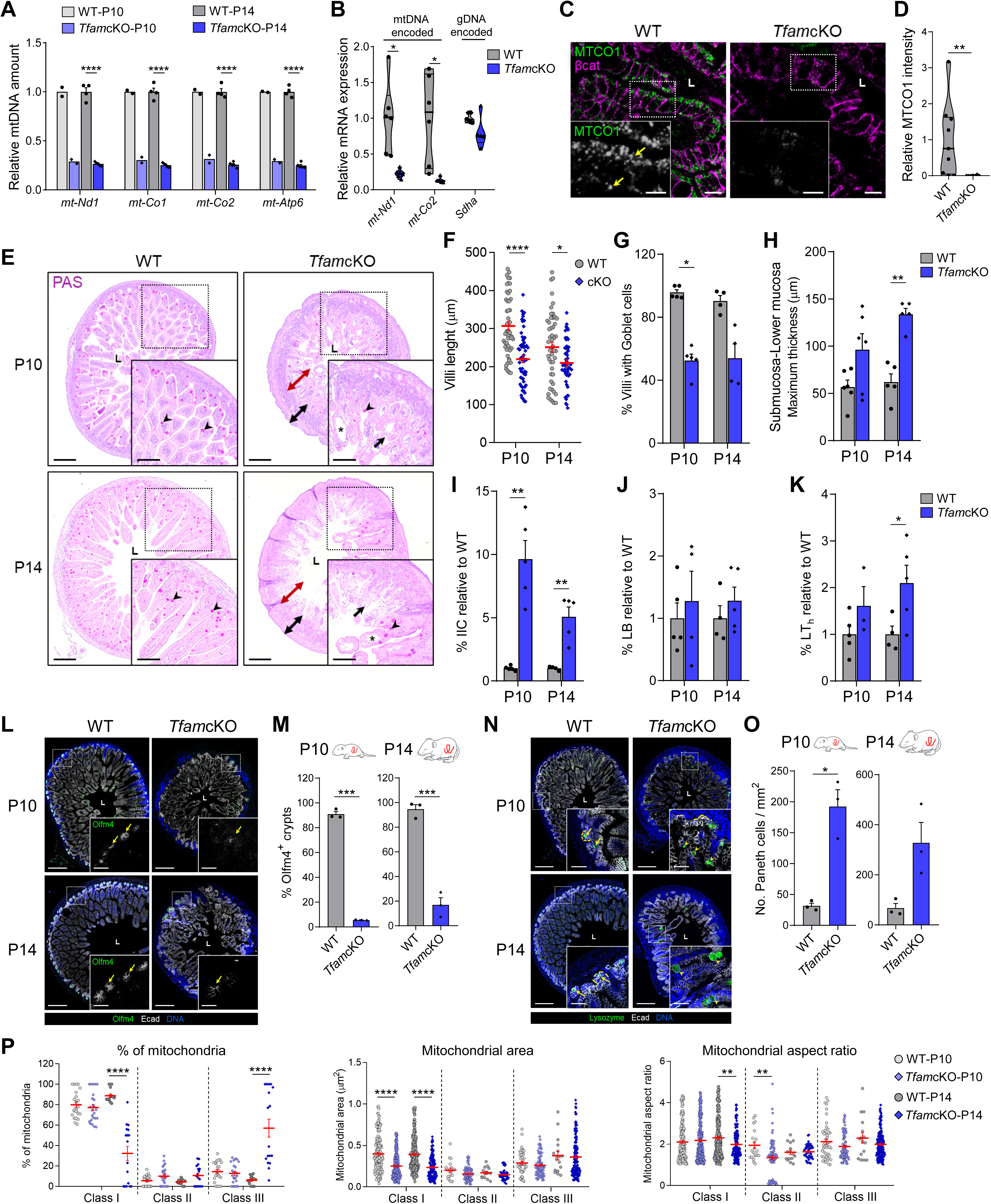
Validation and Intestinal Phenotype of TfamcKO Neonatal Mice. **Related to Figure 6** **(A)** Relative mitochondrial DNA (mtDNA) levels of *Nd1*, *Co1*, *Co2*, and *Atp6* in ileal epithelial cells from WT and *Tfam*cKO mice at P10 (n = 2) and P14 (n = 4), measured by qPCR. **(B)** RT-qPCR analysis of mitochondrial genes *mt-Nd1*, *mt-Co2*, and *Sdha* in ileal epithelial cells from WT and *Tfam*cKO mice at P10 (n = 5–6). **(C)** Confocal images of ileal sections from WT (n = 3) and *Tfam*cKO (n = 2) mice at P14, labelled with anti-MTCO1 (green) and anti-β-catenin (magenta). Arrows point to mitochondria in WT LREs. **(D)** Violin plot shows MTCO1 intensity (integrated density) from (C), normalised to epithelial area. Dots represent individual images (n = 9,7; 3–4 images per mouse). **(E)** PAS and haematoxylin staining of ileal sections from WT and *Tfam*cKO mice at P10 and P14 (n = 4–6) shows inflamed and disorganized epithelium in *Tfam*cKO mice. Phenotypes include shortened villi (double red arrows), thickened submucosa and lower mucosa (double black arrows), oedema (asterisks), reduced PAS+ goblet cells (arrowheads), and abnormal vacuoles in enterocytes (arrows). Scale bars: 200 μm (main); 100 μm (insets). **(F–H)** Quantification of villi length (F), percentage of villi with goblet cells (G), and the maximum thickness of submucosa and lower mucosa (H) from (E). In (F), each dot represents a villus (n = 50,60,50,50; 10 villi per mouse). For (G–H), n = 4–6 mice. **(I–K)** Flow cytometry analysis of innate immune cells (IIC, I), B lymphocytes (LB, J), and T-helper cells (LT_h_, K) percentages in the ileal of WT and *Tfam*cKO mice at P10 and P14 (n = 3–5). See Figure S3 for gating strategy. **(L–O)** Confocal imaging and quantification of stem and Paneth cells in WT and *Tfam*cKO ileal at P10 and P14: • (**L**) Olfm4 (green) labels crypt stem cells; arrows point to Olfm4⁺ crypts. • (**M**) Quantification of Olfm4⁺ crypts (30-100 crypts/mouse). • (**N**) Lysozyme (green) marks Paneth cells; arrows show normal crypt localization, arrowheads indicate mislocalised Paneth cells in villi. • (**O**) Quantification Paneth cells/mm² of epithelial area. For (L, N), sections are also stained with anti-Ecad (white) and DAPI (blue). Scale bars: 200 μm (main); 50 μm (insets). For all panels, n = 3 mice. (P) Distribution of mitochondrial integrity classes in *Tfam*cKO neonatal enterocytes. Quantification of the percentage, area and aspect ratio (long axis / short axis) of each mitochondria class in Figure 6E, as defined in Figure 4B. In percentage graph, dots represent individual enterocytes; in area and aspect ratio graphs, dots are individual mitochondria. Sample sizes are indicated in Table S1. In (C, E, L, N), dashed boxes indicate magnified areas; L, lumen. In (A, B, D, I–K), data relative to WT values. In (A, G–K, M, O), graphs represent mean + SEM and dots represent individual mice. In (B, D), violin plots show the median (thick line) and IQR (thin lines). In (F, P), dot plots show mean ± SEM (red lines). Statistical analyses: Student’s t test (A, B, I–K, M, O), Mann–Whitney test (D, P), two-way ANOVA (F, H), Kruskal–Wallis test (G). In (F–H), only statistically significant comparisons between WT and *Tfam*cKO are indicated within each time group. *p < 0.05, **p < 0.01, ***p < 0.001, ****p < 0.0001.

**Figure S7.**
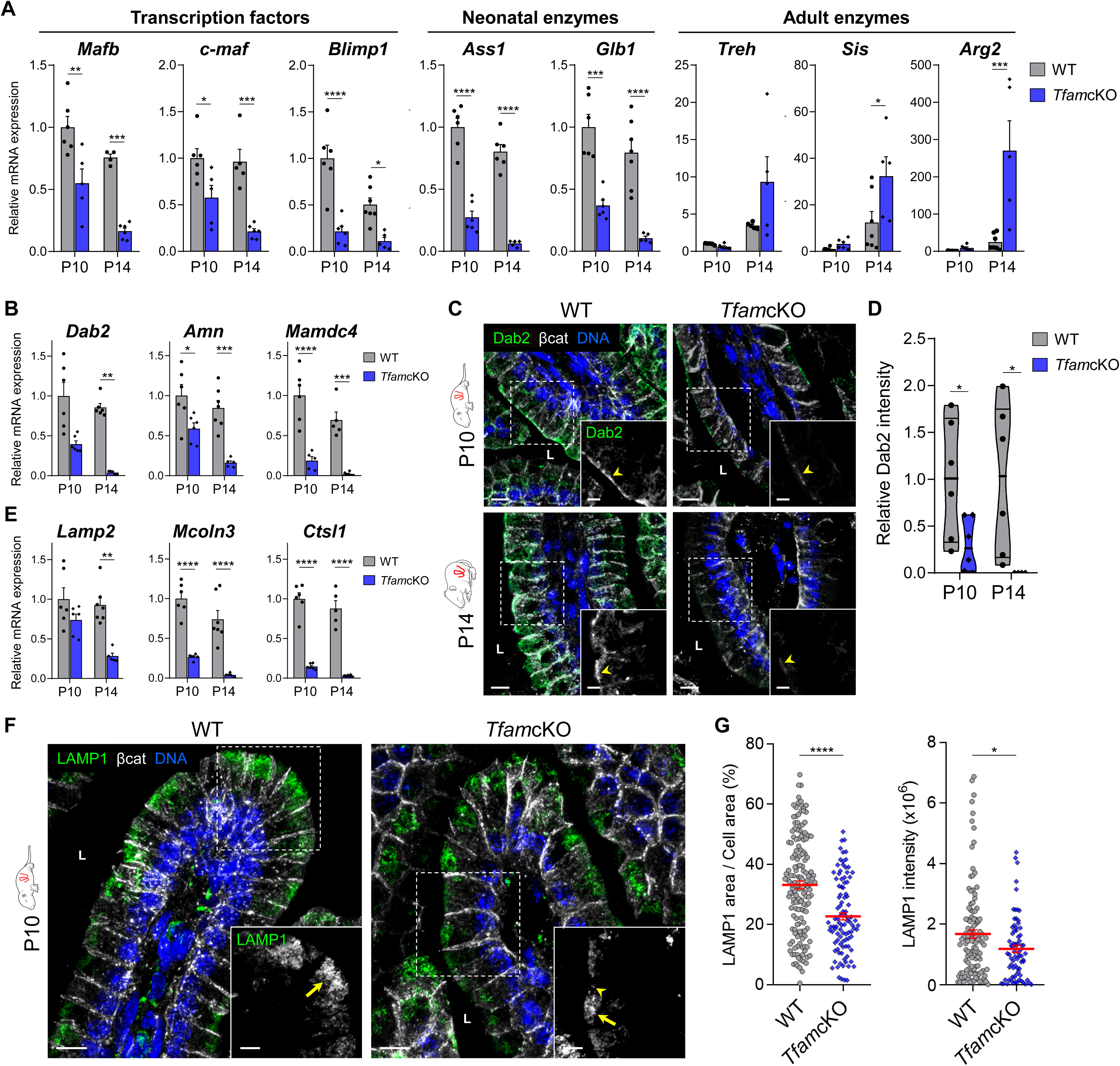
TfamcKO Neonatal Enterocytes Exhibit a Decreased Endolysosomal System and Undergo an Early Metabolic Switch. **Related to Figure 6** **(A)** RT-qPCR analysis of transcription factor (*Mafb*, *c-maf*, *Blimp1*), neonatal (*Ass1*, *Glb1*) and adult (*Treh*, *Sis*, *Arg2*) enzyme genes in ileal epithelial cells from WT and *Tfam*cKO mice at P10 and P14. **(B)** RT-qPCR analysis of *Dab2*, *Amn*, and *Mamdc4* genes in ileal epithelial cells from WT and *Tfam*cKO mice at P10 and P14. **(C)** Confocal images of ileal sections from WT and *Tfam*cKO mice at P10 (n = 3) and P14 (n = 2–3), labelled with anti-Dab2 (green), anti-β-catenin (white), and DAPI (blue). Arrowheads indicate apical Dab2 localisation. **(D)** Violin plot shows apical Dab2 intensity (integrated density) from (C), normalised to epithelial area and relative to WT values. Dots represent individual images (n = 6, 6, 6, 4; 2 images per mouse). **(E)** RT-qPCR analysis of *Lamp2*, *Mcoln3*, and *Ctsl1* in ileal epithelial cells from WT and *Tfam*cKO mice at P10 and P14. **(F)** Confocal images of ileal sections from WT and *Tfam*cKO mice at P10 (n = 3), labelled with anti-LAMP1 (green), anti-β-catenin (white), and DAPI (blue). Arrows indicate endolysosomal system in WT cells; arrowheads highlight the reduced lysosomal vesicles in *Tfam*cKO enterocytes. **(G)** Quantification of lysosomal occupation and content in (E). Dot plots show the LAMP1⁺ area as a percentage of cell area (n = 157,99) and LAMP1 intensity (integrated density) per enterocyte (n = 117,74). Dots represent individual enterocytes. In (C, F), dashed boxes indicate magnified areas; L, lumen. Scale bars: 10 μm (main); 5 μm (insets). In (A, B, E), data relative to WT-P10 values, dots represent individual mice (n = 4–7), and graphs represent mean + SEM. In (D), violin plot shows the median (thick line) and IQR (thin lines). In (G), dot plots depict mean ± SEM (red lines). Statistical analyses: Kruskal–Wallis test (A [*Treh*], B [*Dab2*], E [*Lamp2*]), two-way ANOVA (A, B, E; except *Treh*, *Dab2*, *Lamp2*), Student’s t test (D), Mann–Whitney test (G). In (A, B, E), only statistically significant comparisons between WT and *Tfam*cKO are indicated within each time group. *p < 0.05, **p < 0.01, ***p < 0.001, ****p < 0.0001.

**Table S1.** Sample sizes in TEM graphs. **Related to Figures 4, 6 and S6** Values in the number (No.) and percentage columns correspond to enterocytes; in the area and aspect ratio columns, values correspond to individual mitochondria. For more detailed information on sample size of the analyses, see the TEM section in the Methods.

| | Class | No. mitochondria / $\mu\text{m}^2$ | % of mitochondria | Mitochondrial area | Mitochondrial aspect ratio |
| --- | --- | --- | --- | --- | --- |
| Figure 4C | I | 11, 10 | 11, 10 | 239, 271 | 239, 276 |
|  | II | 11, 10 | 11, 10 | 13, 32 | 12, 28 |
|  | III | 11, 9 | 11, 10 | 14, 12 | 18, 12 |
| Figures 6F and S6P | I | 23, 24, 13, 17 | 23, 24, 13, 17 | 184, 321, 259, 128 | 169, 329, 251, 130 |
|  | II | 23, 24, 13, 17 | 23, 24, 12, 17 | 19, 73, 14, 45 | 21, 74, 15, 44 |
|  | III | 23, 24, 13, 17 | 23, 24, 13, 17 | 45, 78, 17, 134 | 47, 78, 16, 149 |

**Table S2.**
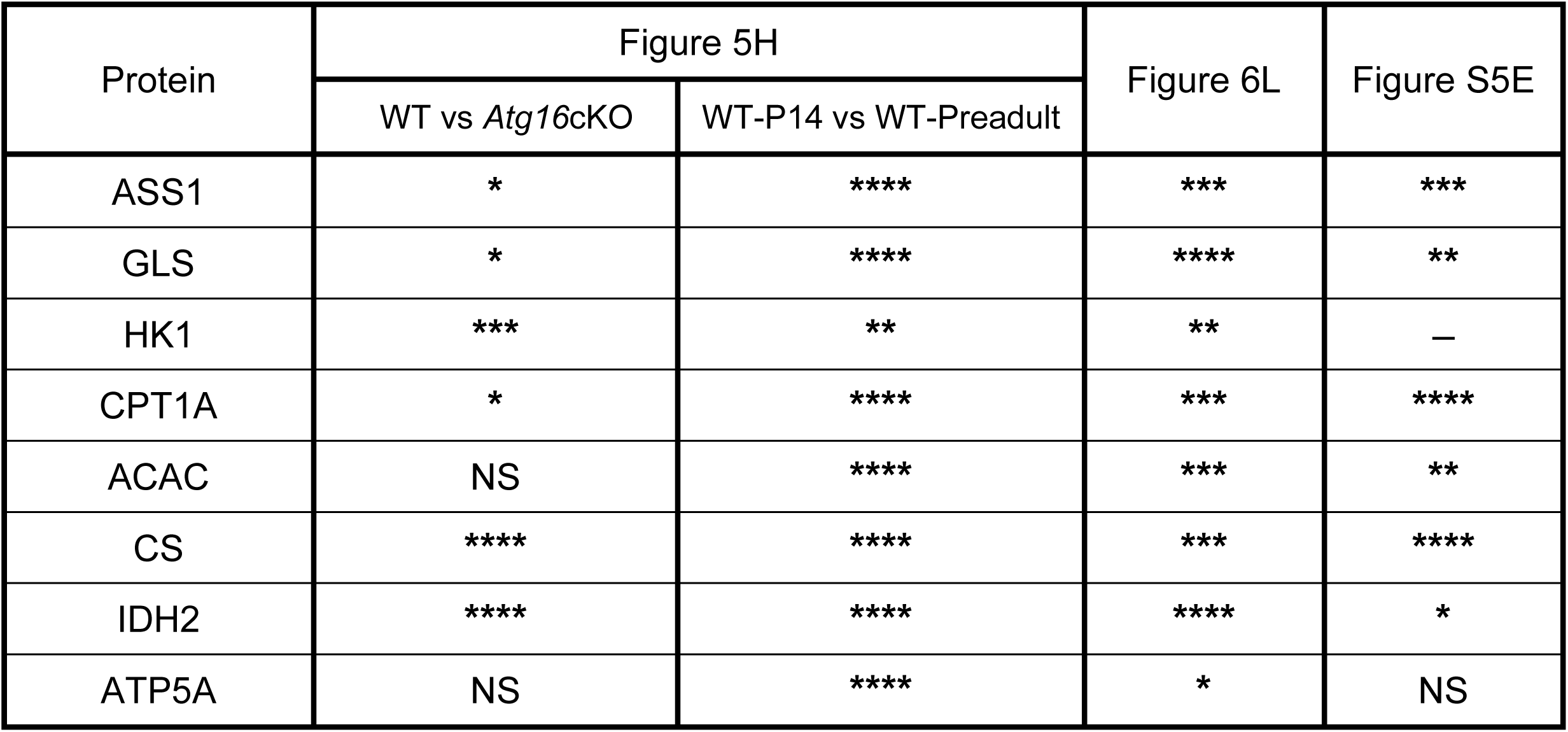
Statistical Significance of Met-Flow Experiments. **Related to Figures 5, 6 and S5** For Figure 5H, statistical results are shown only for comparisons between WT and the two other groups. No statistical test was applied for HK1 of Figure S5E (n = 2). *p < 0.05, **p < 0.01, ***p < 0.001, ****p < 0.0001. NS, not significant.

## METHODS

### KEY RESOURCES TABLE

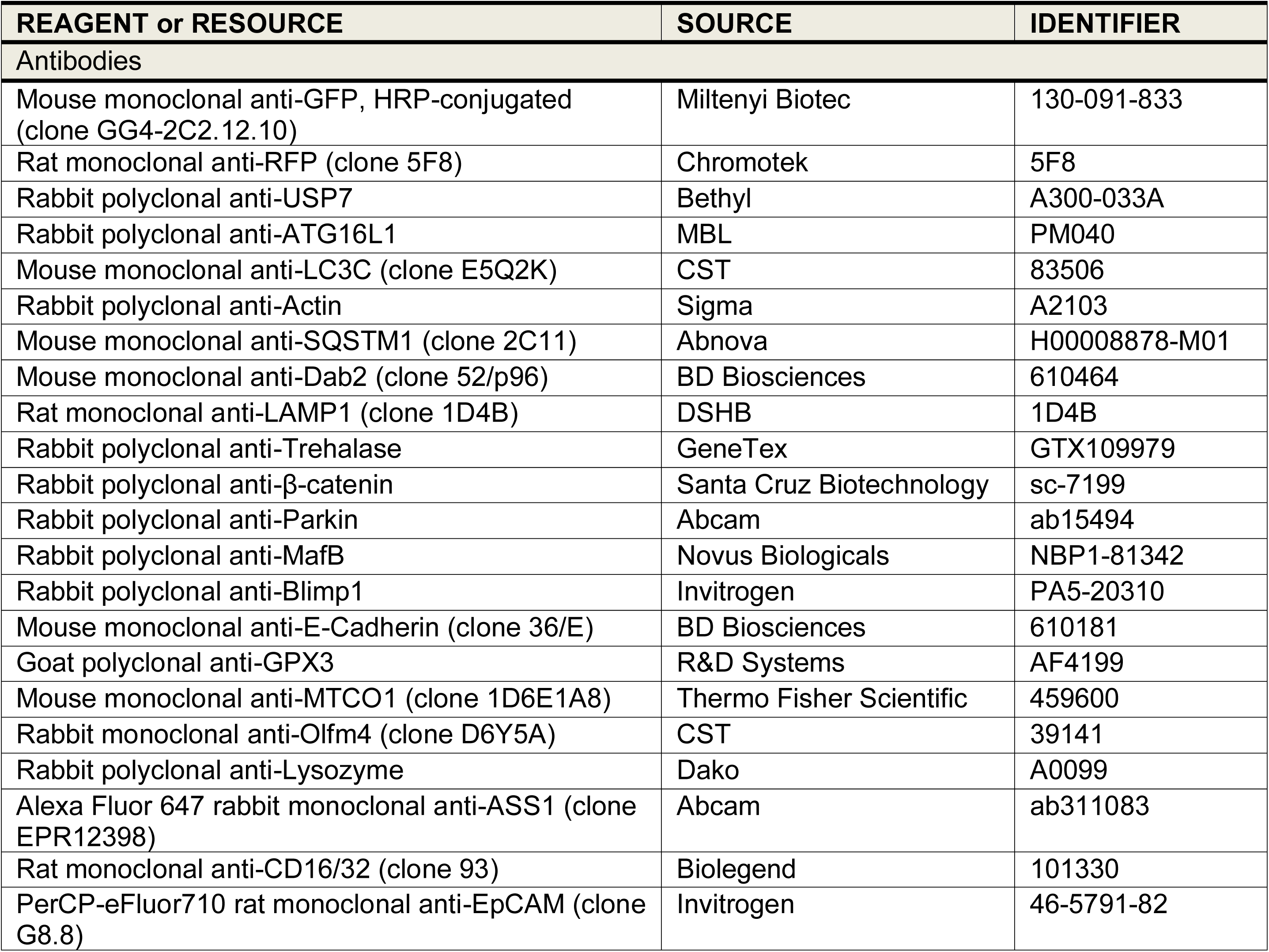
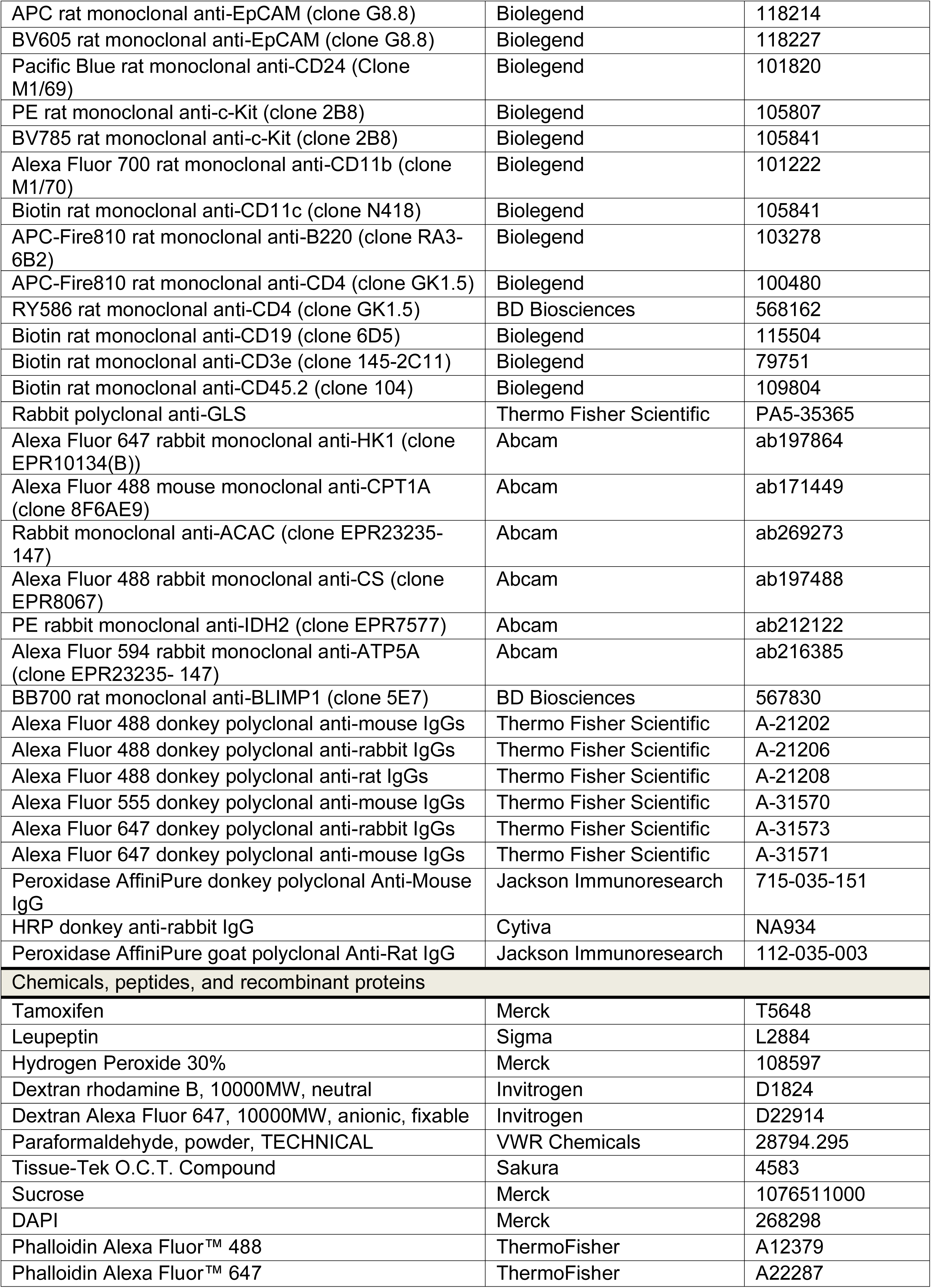
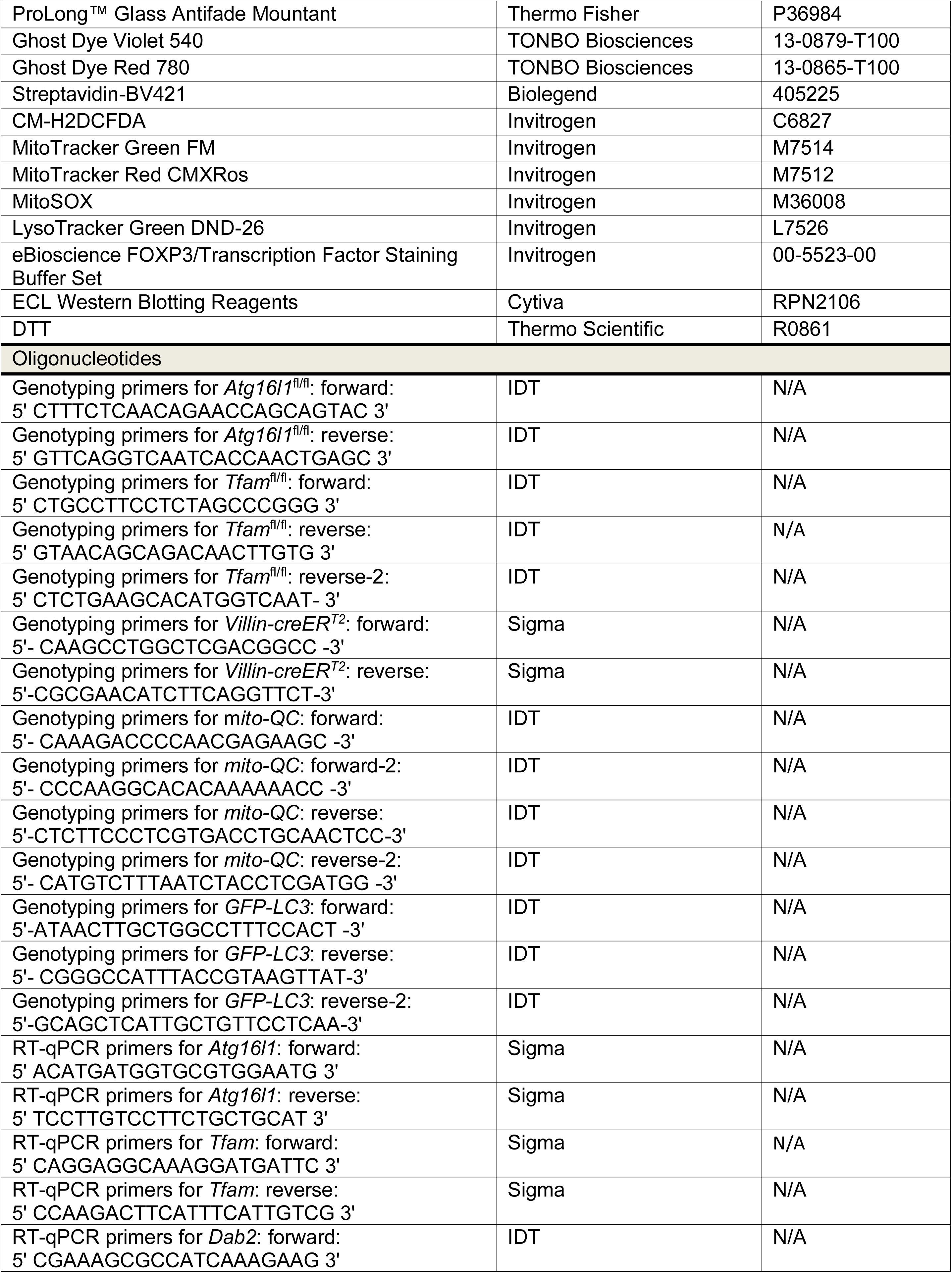
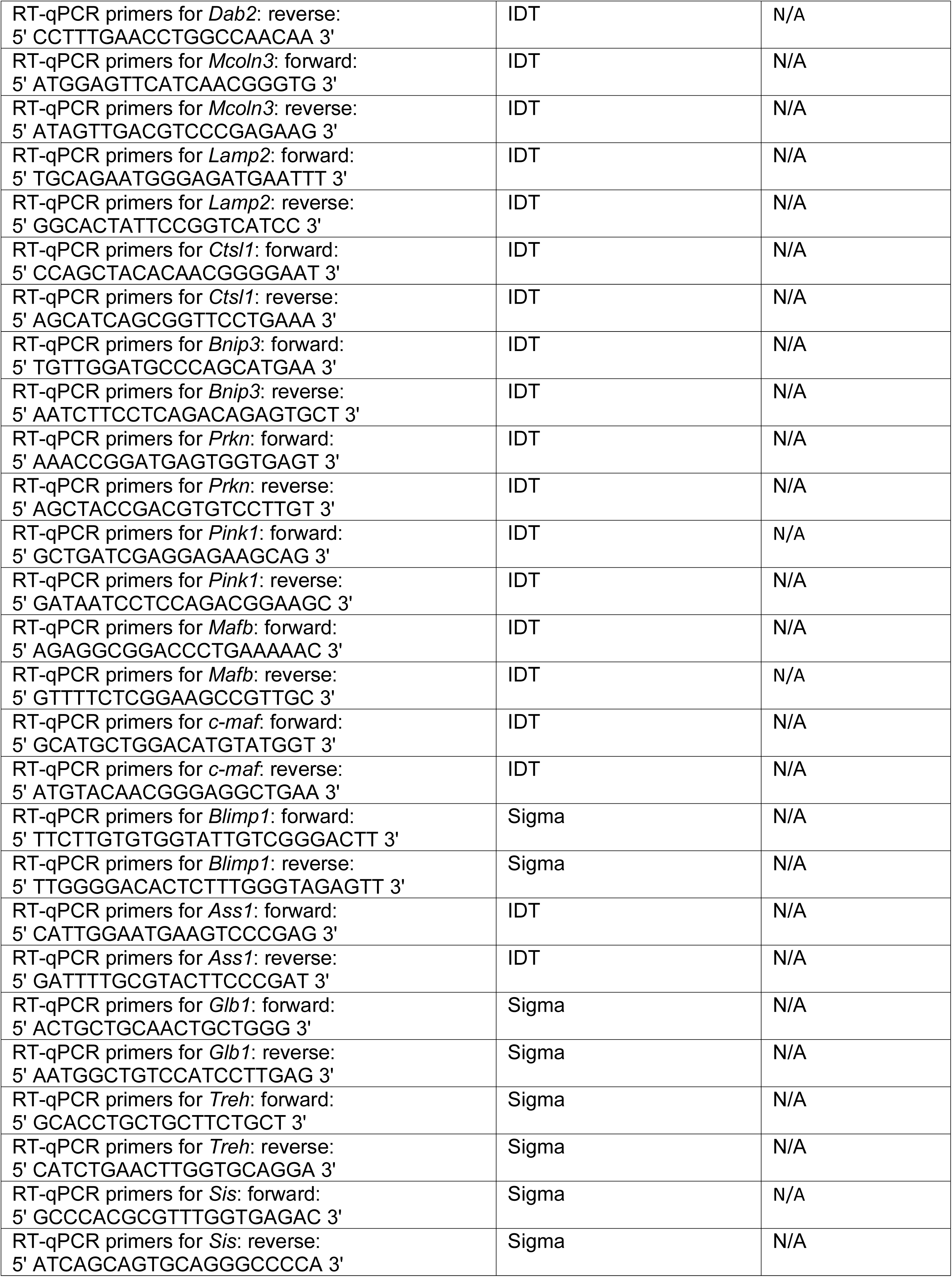
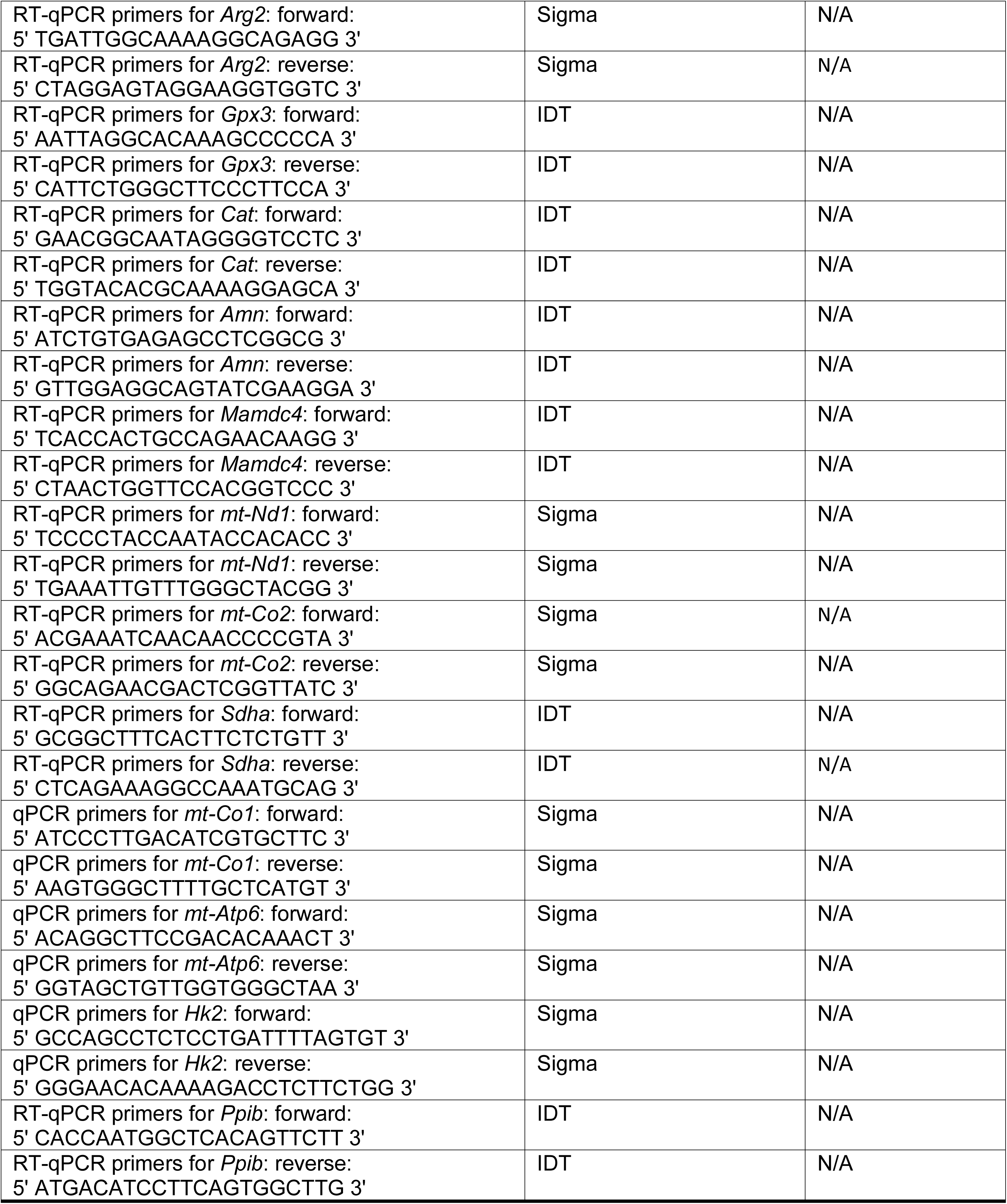

#### Mice

*Atg16l1*^fl/fl33^, *Tfam*^fl/fl^ ^76^, and GFP-LC3^26^ mice were kindly provided by N. Martínez-Martín (Centro de Biología Molecular “Severo Ochoa”, CBM, CSIC-UAM, Madrid, Spain). The *mito*-QC mouse line^42^ was obtained from P. Boya (Centro de Investigaciones Biológicas Margarita Salas, CSIC, Madrid, Spain; University of Fribourg, Switzerland). *Villin-creER^T^*^2^ mice^34^ were provided by N. Djouder (Spanish National Cancer Research Centre, CNIO, Madrid, Spain). *Atg16l1* ^fl/fl^, *Tfam* ^fl/fl^ mice were crossed at the CBM animal facility with *Villin-creER^T^*^2^ mice to generate intestinal epithelial-specific conditional knockout (cKO) mice: *Atg16*cKO and *Tfam*cKO, respectively. In turn, *Atg16*cKO mice were crossed with *mito*-QC mice to establish *the mQ*-*Atg16*cKO mouse line. Tamoxifen (0.12–0.5 mg in corn oil containing 10% ethanol) was administered subcutaneously at postnatal days 1 and 2 to pups from *Atg16*cKO, *mQ*-*Atg16*cKO, and *Tfam*cKO litters, following a protocol similar to that described by Gong et al.^77^. Cre-positive animals were classified as KO, and Cre-negative mice as WT. To confirm the absence of spontaneous recombination in WT mice, expression levels of Atg16l1 and Tfam were assessed by Western blot and RT-qPCR. Mice were genotyped by PCR using DNA extracted from tail biopsies following standard protocols. Genotyping primers are listed in the Key Resources Table. Neonatal mice were humanely sacrificed by decapitation; adult mice were euthanized via CO₂ inhalation. *Tfam*cKO neonates were sacrificed after two consecutive days without weight gain or upon observing approximately a 10% weight loss, in accordance with humane endpoint criteria. No test was performed to determine the influence of sex, as no apparent differences were found. Mouse colonies were maintained at CBM under specific pathogen-free conditions and on a C57BL/6 genetic background. Mice were housed four/five per cage with food and water available ad libitum and maintained in a temperature-controlled environment on a 12 h–12 h light–dark cycle with light onset at 08:00. Animal experiments were performed under protocols approved by the CBM and Comunidad Autónoma de Madrid ethical committees under the following protocol approval number (PROEX 307.2/21). All animal procedures conformed to EU Directive 86/609/EEC and Recommendation 2007/526/EC regarding the protection of animals used for experimental and other scientific purposes, enforced in Spanish law under Real Decreto 1201/2005.

#### mCherry / fluorescent-dextran uptake assay

Mice from GFP-LC3, *Atg16*cKO, and *Tfam*cKO lines were fasted for 1 hour and orally gavaged with either mCherry or 10 kDa fluorescent-dextran (0.3–1 mg/ml in PBS; 10 µl/g body weight). Following gavage, mice were sacrificed either 3 or 16 hours later, adapted from the protocol reported by Remis et al.^12^. To test the endocytic function in neonatal enterocytes, WT mice at postnatal day 7 (P7) were sequentially gavaged with two distinct 10 kDa fluorescent-dextrans (0.7 mg/ml in PBS; 10 µl/g body weight), following a modified version of the protocol established in mouse embryos by Kawamura et al.^78^. After a 1-hour fasting period, pups were first gavaged with Alexa Fluor 647-dextran and then returned to their mother. Three hours later, they were given rhodamine B–dextran again. Pups were subsequently sacrificed either 1 or 5 hours after the second gavage, and ileal enterocytes were analysed by confocal microscopy to assess dextran uptake and intracellular localization.

#### Leupeptin and hydrogen peroxide treatment

To inhibit lysosomal proteases and block autophagy, mice were injected subcutaneously with 40 mg/Kg leupeptin (Sigma) dissolved in PBS and sacrificed after 16h^43^. To assess the effect of ROS on intestinal epithelial maturation, mice were injected intraperitoneally with hydrogen peroxide (5.1 µmol/g body weight; Merck) once daily from P5 to P12 (day of sacrifice), following a protocol adapted from Liang et al.^60^.

#### Histology and Immunofluorescence

Portions of the distal small intestine from mice were fixed in 4% PFA for 16 hours at 4 °C, embedded in paraffin resin, and cut transversely into 5-µm sections. Some samples were stained with hematoxylin-eosin (HE&E; Sigma and Merck, respectively), while others underwent Periodic Acid-Schiff (PAS) staining. Additionally, some samples were used for immunofluorescence. For immunofluorescence, sections were deparaffinized using xylene and then rehydrated through a series of decreasing ethanol concentrations (100%, 96%, 70%, 50%). For antigen retrieval, tissue was boiled at 100 °C on a heat block in sodium citrate buffer (10 mM Sodium citrate, 0.05% Tween 20, pH 6.0) for 20 min. Next, the slides were washed with water and PBS containing calcium and magnesium, and then incubated for 1 hour at room temperature with 3% bovine serum albumin (BSA) as a blocking solution. For O.C.T. embedded sections, portions of the distal small intestine were previously fixed in 4% PFA for 16 hours at 4 °C, incubated in 30% sucrose in PBS for 48-72 hours at 4 °C, and then embedded in O.C.T. compound and stored at -80 °C until use. O.C.T. blocks were cut transversely into 15 µm sections using Superfrost Plus Adhesion Microscope Slides (Epredia) and stored at -80 °C. For immunofluorescence, sections were dried at room temperature for 30 minutes, washed with 0.1% Tween 20, and blocked for 1 hour at room temperature with BGT buffer (0.3% BSA, 0.75% Glycine, 0.3% Triton X-100 in PBS). For both paraffin-embedded and O.C.T.-embedded samples, the primary antibody was added and incubated overnight at 4 °C in the respective blocking buffer. After washing with PBS, the secondary antibody was incubated with DAPI for 1h at RT. Lastly, slides were assembled with coverslips with Prolong Glass Antifade (Thermo Fisher). Antibodies used are specified in the key resources table.

#### Image Acquisition and Analysis

HE&E and PAS staining images were acquired using an Axio Imager M1 vertical wide-field microscope, coupled with a Leica DMC6200 CMOS Colour camera. The objective used was 10X/0.3 Air EC Plan-Neo. LASXv4.13 software was used to stitch the tiles into an image. The analysis of goblet cells, villi length, and submucosa–lower mucosa thickness was performed using Fiji (ImageJ 1.54f; https://fiji.sc/) (Wayne Rasband, NIH, USA)^79^. Lower mucosa was defined as the sum of the *muscularis mucosae*, *lamina propria* and epithelial compartment of the crypts. For villi length analysis, the 10 longest villi per section were chosen. One section per mouse was analysed from 3–6 mice per genotype. For immunofluorescence, images were acquired using a Zeiss LSM 900 confocal microscope mounted on an Axio Imager Z2 upright stand, with sequential acquisition and bidirectional scanning. For individual fields, a 63×/1.4 NA oil-immersion Plan-Apochromat objective was used, with a voxel size of 50 × 50 × 520 nm. For tile scans, images were acquired using a 20×/0.8 NA air Plan-Apochromat objective, with a voxel size of 0.624 × 0.624 × 1.9 µm, and stitched using ZEN Blue software version 3.4 (Zeiss; https://www.zeiss.com/zen). For the analysis of Paneth cells and Olfm4⁺ crypts in the ileum of *Tfam*cKO mice, all cells and all crypts were quantified per section (one section per mouse; 3 mice per genotype). For intensity analysis, all images were acquired with 16-bit resolution and under the same conditions. An average intensity projection was made, and the endothelial cells (lacteal vessel) were cropped. The same threshold was applied to all images to select positive areas. Images were analysed using CellProfiler 4.2.8 (https://cellprofiler.org/)^80^ and Fiji (ImageJ 1.54f)^79^. Intensity values correspond to the integrated density, normalized to the epithelial nuclear area (defined by DAPI in epithelial cells) for BLIMP1 and MAFB, or to the epithelial area (delimited by β-catenin, E-cadherin, or F-actin) for all other markers, except for LAMP1 (in both KO models) and Rd-dextran (only in *Atg16*cKO mice), which were not normalized. For LAMP1 intensity and area analysis in *Tfam*cKO mice, images were deconvolved using Huygens Professional software (version 24.10; Scientific Volume Imaging, https://svi.nl/Huygens-Professional). For the analysis of Rd-dextran intensity and LAMP1 area and intensity in *Atg16*cKO mice, 25–40 and 26–36 enterocytes per condition were analysed, respectively. For LAMP1 area and intensity analysis in *Tfam*cKO mice, a minimum of 30 enterocytes per mouse were quantified, across 3 mice per group (99–167 enterocytes per group). For all other intensity analyses, 2–5 random images per mouse were acquired from comparable villi, each containing approximately 50 cells, with 2–5 mice per group.

#### Transmission Electron Microscopy

Samples were processed by the Electron Microscopy Unit at CBM. Distal small intestine portions were fixed using 4% PFA (paraformaldehyde) and 2% glutaraldehyde in phosphate buffer 0.1M, pH 7.4, for 2 hours at room temperature. Then they were washed 3 times for 10 minutes with phosphate buffer. After fixation, the material was post-fixed in 1% osmium tetroxide and potassium ferricyanide for 1 hour at 4 °C, then embedded in tannic acid for 1 minute and block-stained with 2% uranyl acetate for 1 hour in the dark. Next, they were dehydrated in crescent concentrations of ethanol (30%, 50%, 70%, 95%, 3x100%; 10 minutes each) and infiltrated in propylene oxide for 45 minutes and Epon 100% for 1 hour. Finally, the samples were introduced into BEEM capsules (ESBE Scientific). Samples were observed using a JEM1400 Flash Transmission Electron Microscope (JEOL) equipped with a CMOS OneView camera (Gatan). Fiji (ImageJ 1.54f)^79^ was used for image analysis. Mitochondria were classified into three categories, as in Moschandrea et al.^44^: class I (normal), characterized by mitochondria with narrow and well-defined cristae; class II (partly affected), with mitochondria exhibiting undefined cristae; and class III (damaged), including mitochondria with wide, short, and less electron-dense cristae, or those with ruptured membranes or internal inclusions. For the analysis of mitochondrial status in *Atg16*cKO mice, 291–340 mitochondria were analysed from 10– 11 enterocytes (3 mice per genotype). For *Tfam*cKO mice, in the analysis of number and percentage of mitochondrial classes, 444–1040 mitochondria from 13–24 enterocytes (3–4 mice per group) were used; for area and aspect ratio analysis, 261–520 mitochondria from 11–17 enterocytes (2–4 mice per group) were analysed. The graphs show data after outlier exclusion.

#### Isolation of IEC

Mice were euthanized, and the abdominal cavity was opened to expose the intestine. The region of interest (proximal or distal intestine) was dissected, removed, and rinsed in PBS. The tissue was cut transversely into ∼5 mm fragments, which were then opened longitudinally and incubated in epithelial dissociation buffer (1 mM DTT, 1 mM EDTA, 2% FBS in PBS) at 37°C with agitation for 20 minutes. After incubation, the samples were centrifuged at 1200 rpm for 5 minutes. The supernatant was discarded, and the pellet was resuspended in 10 mL of DMEM supplemented with 10% FBS at 4 °C. The suspension was passed through 70 μm cell strainers (Falcon) and then centrifuged again at 1,200 rpm for 5 minutes. The resulting pellet was resuspended in FACS buffer (1 mM EDTA, 2% FBS in PBS) at 4 °C and centrifuged again under the same conditions. The final pellet containing epithelial cells was used for western blotting, flow cytometry, and RT-qPCR analysis.

#### Immunoblotting

IECs pellets were lysed with lysis buffer (50 mM Tris-HCl, 150 mM NaCl, 1% NP-40, 50 mM NaF, 0.1% SDS, 1 mM Na_3_VO_4_, 30 mM β-glycerophosphate, 5 mM EDTA, protease inhibitors) and centrifuged for 10 minutes at 4 °C, 15,000 rpm. Supernatants were collected and quantified using the Bradford protein assay (Biorad Protein Assay, Biorad). 25 µg of protein was loaded in 10 or 12% acrylamide gels for SDS-PAGE. After electrophoresis, proteins were transferred to an Immobilon-P transfer membrane (Millipore), blocked for 1 hour, and incubated overnight with primary antibodies at 4 °C. After washing with 0.05% Tween 20, they were incubated with HRP-conjugated secondary antibodies for 1 hour at room temperature. ECL western blotting reagents (Cytiva) were used for the chemiluminescence reaction. Bands were analysed using Fiji (ImageJ 1.54f)^79^. Antibodies used are specified in the key resources table.

#### Flow Cytometry

Disaggregated epithelial cells were plated into a 96-well plate and incubated with FACS buffer with anti-CD16/CD32 Fc block and viability markers (Ghost Dye) for 10 min at room temperature. After centrifugation, cells were incubated for 20 min at 4°C with the appropriate combination of the antibodies in FACS buffer to discern between the different epithelial and immune populations. Following protocols adapted from immunometabolic ^45,54^ and intestinal studies^36^, we assessed mitochondrial mass and membrane potential, cellular and mitochondrial ROS levels, and lysosomal content labelling cells with MitoTracker Green FM (50 nM), MitoTracker Red CMXRos (100 nM), CM-H2DCFDA (2.5 µM), MitoSOX (1 µM), and Lysotracker Green DND-26 (100 nM), respectively. Probes were incubated for 15 min at room temperature and 15 min at 37 °C in DMEM with 5% FBS (Lysotracker), without FBS (MitoTracker and CM-H2DCFDA), or in HBSS (MitoSOX). Antibodies used are specified in the key resources table. Data were acquired using a Cytek Aurora full-spectrum cytometer at the CBM, and analysed with FlowJo software (versions 10 and 10.8.1; BD Biosciences). See Figure S3 for details on the gating strategies.

#### Met-flow cytometry staining

Based on recent works on immune cell metabolism^45,50^, we selected eight key metabolic enzymes to analyse their expression by flow cytometry in our epithelial cells: ASS1, GLS, HK1, CPT1A, ACAC, CS, IDH2, and ATP5A. After the disaggregated epithelial cells were incubated with surface antibodies according to the flow cytometry protocol, they were fixed and permeabilized using the eBioscience Foxp3/Transcription Factor Staining Buffer Set (Invitrogen). Next, they were washed in the provided wash buffer and blocked with 20% FBS in wash buffer and incubated for 2h at room temperature with the appropriate antibodies to detect the rate-limiting enzymes from different metabolic pathways. Lastly, they were rewashed with wash buffer. Antibodies used are specified in the key resources table. Data were acquired using a Cytek Aurora full-spectrum cytometer at the CBM, and analysed with FlowJo software (versions 10 and 10.8.1; BD Biosciences).

#### RNA Sequencing and Bioinformatic Analyses

The FACS gating strategy used to isolate LREs and PECs for RNA sequencing (RNA-seq) is shown in Figure S4A. Cell sorting was performed using a FACSAria Fusion cytometer (BD Biosciences). Each population was collected in biological triplicate samples of 300,000–500,000 cells. For LREs, each replicate consisted of pooled cells from multiple neonatal litters; for PECs, each replicate corresponded to a single mouse at postnatal day 42 (P42). Cells were collected in RLT buffer (Qiagen), and total RNA was extracted using the RNeasy Plus Micro Kit (Qiagen). Samples with an RNA integrity number (RIN) >7.0 were selected for RNA-seq. Ultra-low input poly(A)-selected libraries (single-end) were prepared in triplicate and sequenced using the Illumina HiSeq 2500 platform (50 bp, single-end reads) at the Centro de Análisis Genómico (CNAG-CRG, Barcelona, Spain). Reads were aligned to the Mus musculus reference genome (mm39) using HISAT2 (v2.1.0), and gene counts were obtained using HTSeq-count (v0.11.2). Differential gene expression analysis was performed with DESeq2. Gene Set Enrichment Analysis (GSEA)^81^ was conducted using the clusterProfiler ^82^ and enrichplot packages within the R environment. Predefined gene sets from the Gene Ontology (GO) database were assessed for statistically significant enrichment based on a ranked list of gene expression values. The full gene dataset, annotated with ENSEMBL identifiers and their associated statistical metrics (stat), was utilized as input for the GSEA. Enrichment was quantified by normalized enrichment scores (NES), with significance determined at an adjusted p-value threshold of < 0.05 following Benjamini-Hochberg correction. Over-Representation Analysis (ORA)^37^ was performed in R using the clusterProfiler package. Differentially expressed genes (DEGs) with an adjusted p-value < 0.05 and mapped using ENTREZID identifiers were used. Enriched GO terms and KEGG pathways were also filtered at an adjusted p-value < 0.05 (Benjamini-Hochberg correction). Selected GO and KEGG terms were visualized using bubble plots generated in R with the GOplot package (GOBubble function) ^83^. The biocomputational data analysis was carried out by the Biocomputational Analysis Service (SABio) at the Centro de Biologia Molecular “Severo Ochoa” (CSIC-UAM).

#### RNA Extraction, cDNA Synthesis, and RT-qPCR

Epithelial cells were lysed in 500 µL of TRIzol (Invitrogen) and stored at -80 °C until RNA extraction. RNA was isolated using the RNeasy Micro Kit (Qiagen) according to the manufacturer’s instructions. For cDNA synthesis, 500 ng of RNA was reverse transcribed using the High-Capacity cDNA Reverse Transcription Kit (Applied Biosystems). Gene expression was assessed by quantitative PCR (qPCR) using gene-specific primer pairs (listed in Key Resources Table). Reactions were performed in 384-well plates using either the CFX384 Real-Time System (Bio-Rad) or the ABI 7900HT Fast Real-Time PCR System (Applied Biosystems), with SYBR Green Master Mix (Agilent, Santa Clara, CA). Each well contained 10 ng of cDNA and 2.5 µM of each primer in a final volume of 10 µL. All reactions were run in technical triplicates. Cyclophilin B (*Ppib*) was used as the reference gene for normalisation.

#### mtDNA Quantification

Genomic DNA (gDNA) was extracted using the NZY Tissue gDNA Isolation Kit (NZYTech). Relative mitochondrial DNA (mtDNA) content was determined by qPCR using primers for the following mitochondrial genes: NADH dehydrogenase 1 (*mt-Nd1*), cytochrome c oxidase subunits 1 and 2 (*mt-Co1* and *mt-Co2*), and ATP synthase subunit 6 (*mt-Atp6*). Reactions were performed in 384-well plates using the CFX384 Real-Time System (Bio-Rad) with SYBR Green Master Mix (Agilent, Santa Clara, CA). Each well contained 10 ng of gDNA and 2.5 µM of each primer in a final volume of 10 µL. All reactions were run in technical triplicates. Hexokinase 2 (*HK2*) was used as the reference gene for normalization, with primers listed in the Key Resources Table.

### QUANTIFICATION AND STATISTICAL ANALYSIS

GraphPad software (versions 8.0.1 and 9.5.1) and Microsoft Excel were used for data analysis and plotting. Results shown are either representative or cumulative from at least two independent experiments. Except for gene expression analyses during development, RT-qPCR and WB data were normalized to the average of at least two WT or control mice of their corresponding litter. Sample size, statistical tests, and the meaning of the dots in each graph are specified in figure legends, except for TEM graphs (Figures 4C, 6F and S6O; samples sizes indicated in Table S1). In violin plots, median and interquartile range (IQR) are shown; in dot plots, mean ± SEM; in bar plots with dots, the mean + SEM; and in standard bar plots, the mean + SD. Outliers were identified using the ROUT method (Q = 5%) or Grubbs’ test (α = 0.05) for datasets with n ≥ 6. Normality was assessed using the Shapiro–Wilk test (n < 50) or the Kolmogorov–Smirnov test (n > 50). Statistical significance between two groups was determined using Student’s t-test for parametric data or the Mann–Whitney test for nonparametric data. Welch’s correction was applied when variances between groups were unequal. For comparisons among more than two groups, one-way ANOVA or two-way ANOVA with Tukey’s post hoc test was used for parametric data, and the Kruskal–Wallis test, followed by Dunn’s multiple comparison test, was used for nonparametric data. Survival curves were compared using the Gehan–Breslow–Wilcoxon test. A critical value for significance of p < 0.05 was used throughout the study (*p < 0.05, **p < 0.01, ***p < 0.001, ****p < 0.0001). In Figures 1H, 2A, 2H, S5B–C, S6F–H, S7A, S7B, S7E, and Table S2, only the most relevant statistically significant comparisons are shown; non-significant results are not displayed. In all other figures, statistical significance is indicated only when differences are significant.

